# Vicious circle of amyloid and leptomeningeal macrophages evokes vascular dysfunction in CAA

**DOI:** 10.64898/2026.02.12.705483

**Authors:** Rahadian Yudo Hartantyo, Takahiro Tsuji, Lingnan Hou, Rie Saito, Ayato Yamasaki, Takahiro Kochi, Yutaro Saito, Mari Tada, Dennis Lawrence Cheung, Andrew J Moorhouse, Marco Prinz, Takahiro Masuda, Akiyoshi Kakita, Hiroaki Wake

## Abstract

The subarachnoid space contains leptomeningeal arteries and resident immune cells known as subarachnoid macrophages (SAMs). In Alzheimer’s disease (AD) and related cerebral amyloid angiopathy, amyloid-β (Aβ) frequently accumulates around the leptomeningeal arteries. Nevertheless, the local immune responses and their functional consequences remain poorly understood. Using longitudinal intravital imaging in a mouse model of AD, we tracked Aβ deposition and identified a distinct SAM population that migrates to and phagocytoses these arterial Aβ deposits. SAM recruitment correlated with vascular remodeling, including smooth muscle loss, aneurysm formation, and reduced cerebral perfusion. Transcriptomic profiling revealed a distinct SAM population in AD with upregulated CD39 expression. Pharmacological inhibition of CD39 attenuated arterial Aβ deposition, identifying SAMs as a potential therapeutic target in AD.

## INTRODUCTION

Brain tissue integrity and function are supported by two broad populations of resident immune cells with distinct anatomical locations and transcriptomic signatures: parenchymal microglia and central nervous system (CNS)-associated macrophages (CAMs).^1–3^ As a homeostatic phenotype, the parenchymal microglia actively survey neural components and modulate cerebrovascular reactivity in healthy individuals, although with some transcriptomic, regional, and functional diversity.^4–6^ During brain injury, infection, cancer, or neurodegenerative disease, these homeostatic microglia transform into functionally divergent populations in response to environmental signals specific to the disturbance.^7^ In Alzheimer’s disease (AD), characterized by the development of amyloid-beta (Aβ) plaques, parenchymal microglia transform into disease-associated phenotypes with neuroprotective subpopulations that can phagocytose the Aβ plaques^8^ and subpopulations that can exacerbate the spread of neuroinflammation and the pathological hallmarks of the disease.^9–11^ Single-cell RNA profiling and gene manipulation studies have identified parenchymal microglial signaling pathways activated in the AD landscape, leading to the discovery of novel therapeutic targets that could mitigate cognitive decline in AD.^12–14^

In contrast, only a few studies have assessed the role of CAMs in neurodegenerative diseases, although recent transcriptomic and proteomic studies have helped clarify their molecular profiles.^1,15^ At least six distinct transcriptomic clusters have been suggested, with some exhibiting preferential locations.^16^ CAM populations are found in the perivascular spaces, choroid plexus, cerebrospinal fluid (CSF), and in the dural and subdural meninges.^17,18^ At the interface between the blood–brain barrier (BBB) and the parenchyma, perivascular macrophages were reported to exacerbate synapse phagocytosis in mouse models of AD by secreting a soluble signaling molecule (SPP1) that triggers a microglial phagocytic transformation.^19^ The authors speculated that abnormal Aβ clearance across the BBB may initiate the perivascular macrophage response. Whether other identified CAM populations also respond to signals in their local environments to influence AD pathogenesis remains unclear.

CAMs in the subarachnoid space are particularly interesting in AD. The subarachnoid space is a delicate, CSF-filled region bounded by the arachnoid and pia mater meninges.^17^ The subarachnoid space is richly vascularized and supported by a trabecular scaffold,^20^ with its dense network of arteries essential for nourishing the underlying cerebral cortex and representing the blood–CSF barrier. The subarachnoid space is particularly relevant in AD for two key reasons. First, soluble Aβ peptides within the parenchyma interstitial fluid are thought to be cleared from the brain via a perivascular drainage pathway through the subarachnoid space, before absorption into the lymphatic vessels within the dural meninges.^21^ Soluble components within the cortical parenchymal interstitium readily access the subarachnoid CSF fluid compartment, and pial surface meninges may also be less of a diffusion barrier than previously thought.^22^ Therefore, macrophages and immune cells within the subarachnoid space are strategically positioned as vigilant sentinels, monitoring the brain interstitium and specifically influencing Aβ clearance. Second, the accumulation of Aβ plaques within the subarachnoid, or “leptomeningeal,” arteries is the most common neurovascular comorbidity in AD and a key pathological feature of cerebral amyloid angiopathy (CAA), commonly observed in aged and AD brains.^23,24^ CAA can potentially result in increased cerebral microbleeds and subarachnoid and cerebral hemorrhages.^25^ The importance of understanding the interactions between arterial Aβ deposits, vessel integrity, and local immune processes is exemplified by the occurrence of amyloid-related imaging abnormalities (ARIAs). ARIAs are common adverse events reported in the treatment of AD with anti-Aβ immunotherapy. ARIAs include subarachnoid hematomas arising from antibody-mediated inflammatory reactions and vessel modeling around pre-existing Aβ deposits.^26^

Therefore, we undertook a series of experiments to track the development of Aβ deposits in leptomeningeal vessels in AD model mice. We hypothesized that these deposits would interact directly with subarachnoid macrophages (SAMs).

## RESULTS

### Aβ deposits in the brain and subarachnoid vessels

First, we examined Aβ deposition in the brains of two donors diagnosed with CAA. Occipital cortex blocks were tissue-cleared, immuno-stained for Aβ and vascular smooth muscle, and visualized with light-sheet microscopy (Figure 1A). Abundant Aβ deposition was observed around subarachnoid arteries, typically with a banded pattern, as reported previously ^27^. Three-dimensional reconstruction of the Aβ bands clearly demonstrated their circumferential, coil-like appearance around arteries (Figures 1A inset and S1A-S1D). The quantified volumes of Aβ and vascular smooth muscle were inversely related (Figures 1B and 1C), suggesting that arteries with more extensive Aβ coverage had less smooth muscle.

**Figure 1.**
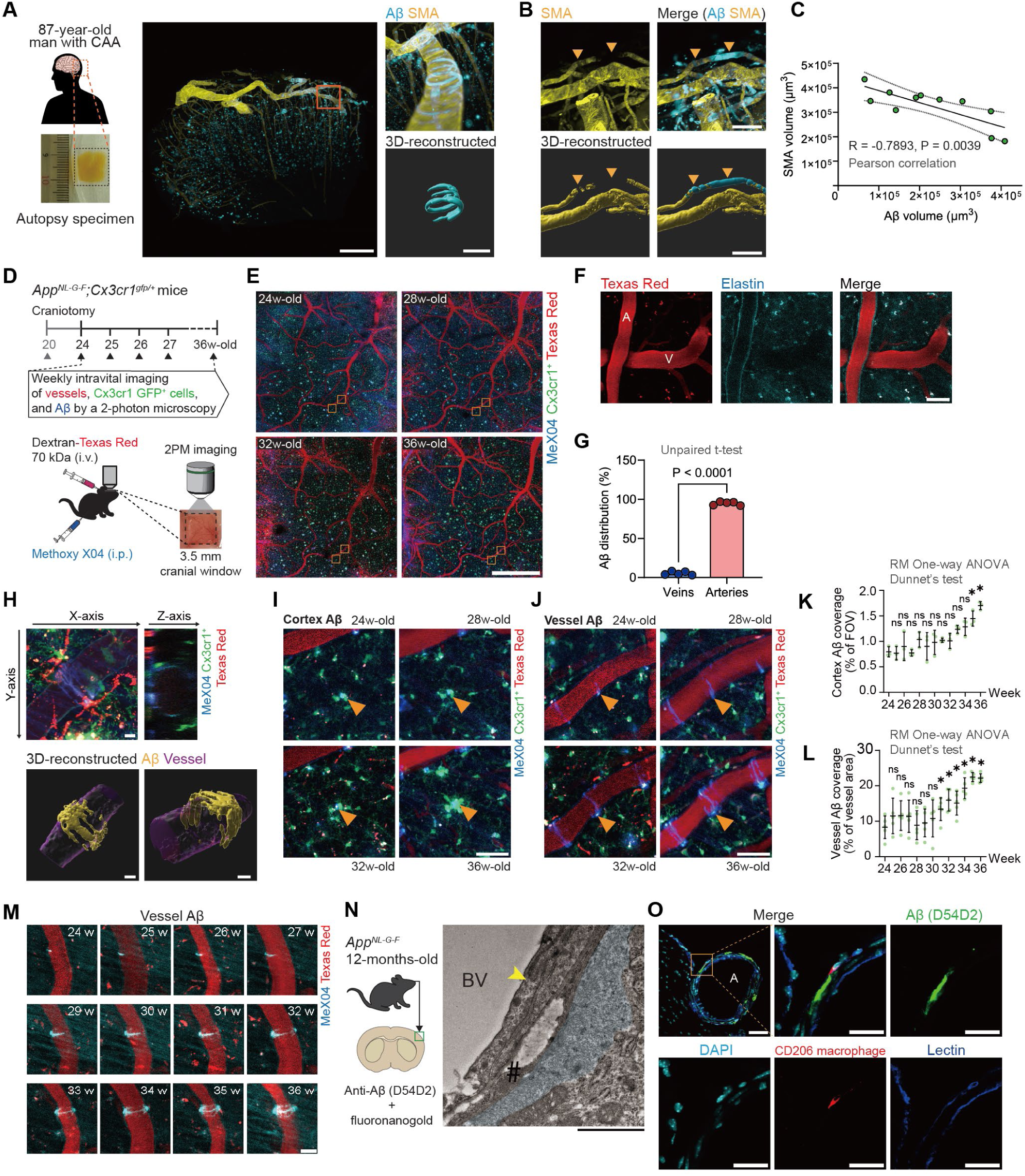
Aβ deposits in the brain and subarachnoid vessels. (**A**) Experimental scheme for the immunostaining and clearing of human brain samples. Typical images of brain samples with cerebral amyloid angiopathy. Note the ring-shaped Aβ deposition formed around the arteries. **(B)** Vascular smooth muscle and Aβ deposition 3D rendering. (**C)** Correlation between Aβ deposition and vascular smooth muscle. (**D)** Experimental scheme. Chronic *in vivo* two-photon imaging was performed in *App^NL-G-F/NL-G-F^Cx3cr1^GFP/+^* mice. The mice were injected intraperitoneally with Methoxy-X04 and intravenously with 70 kDa Texas red dextran to visualize Aβ deposition and vessels, respectively. **(E)** Typical time course images of Aβ deposition, vessels, macrophages, and microglia in *App^NL-G-F/NL-G-F^* mice. **(F)** Autofluorescence with elastin was visualized to distinguish cerebral arteries from veins. **(G)** The summarized graph of the Aβ deposition on arteries. **(H)** An orthogonal view and 3D reconstruction of a typical *in vivo* two-photon image revealing Aβ deposition around the vessel. **(I)** Typical time course images of Aβ deposition in the brain parenchyma. **(J)** Typical time course images of Aβ deposition in the subarachnoid arteries. **(K,L)** The summarized time course graph of the Aβ coverage in the brain parenchyma and subarachnoid arteries. **(M)** Detailed ring-shaped Aβ deposition development around the arteries. **(N)** Electron microscopy of Aβ deposition (*) around the vessel (BV), adjacent to vessel smooth muscle (#) and endothelial cell (yellow arrow). **(O)** Immunostaining images show CD206^+^ macrophages associated with Aβ deposition on vessels (tomato lectin). Data are presented as the mean ± standard deviation (**G**, **K**, **L**). Statistical comparisons were made using unpaired t-tests. N = 5 (**G**, **L**) and 3 (**K**). Scale bar, 1 µm (**N**), 10 µm (**H**–**J**), 25 µm (**M**, **O**-inset), 50 µm (**F**, **O**), 250 µm (**A**-inset, **B**), 500 µm (**A**), 1000 µm (**E**).

We investigated the temporal dynamics, cellular mechanisms, and functional consequences of Aβ deposition on the subarachnoid arteries, using the transgenic *App^NL-G-F^* mouse model of AD ^28^ sequential *in vivo* two-photon imaging from 24 to 36 weeks of age (Figures 1D and 1E). Microglia and macrophages were concurrently visualized by crossing homozygous *App^NL-G-F^* and *Cx3cr1^GFP^* mice. Aβ and blood vessels were identified by injecting methoxy-X04 (intraperitoneal) and 70 kDa Texas Red dextran (intravenous), respectively, before imaging. Arteries were distinguished from veins using the two-photon excitation auto-fluorescence signal of elastin (Figure 1F) ^29^. As observed in the human samples, Aβ accumulated in vessels in aged mice. In the subarachnoid vessels, almost all (98%) Aβ deposits were in the arteries rather than the veins (Figure 1G). Three-dimensional (3D) image-reconstruction demonstrated a coil- or ring-like pattern of Aβ deposition (Figure 1H) resembling that seen in human post-mortem samples.

In addition, Aβ deposition was observed in the cortical parenchyma. Both parenchymal and subarachnoid vessel Aβ accumulation increased over time, with a slightly earlier onset in subarachnoid vessels (from 31 weeks) than in the parenchyma (from 33 weeks; Figures 1I-1L). Arterial Aβ deposition progressed from small spot-like aggregates to ring-like formations over time (Figure 1M; Video S1), supporting previous suggestions that Aβ bands are “seeded” by initial deposits ^27^. Electron microscopy confirmed Aβ deposits around the cerebral arteries, distal to the endothelium, and adjacent to vascular smooth muscle cells (Figure 1N). Furthermore, immunostaining of fixed brain sections identified CD206^+^ immune cells around vessel-associated Aβ deposition (Figure 1O), which resembled macrophages rather than resident parenchymal microglia. In addition, macrophages were closely opposed to the rings of arterial Aβ deposits in the post-mortem human tissue (Figure S1E).

### Macrophage reactivity to subarachnoid Aβ deposits

We conducted time-lapse *in vivo* imaging of SAMs in AD mice at high temporal resolution during Aβ deposition in the leptomeningeal arteries (Figure 2A). Meningeal macrophages, SAMs, and microglia had distinct locations and morphological features, enabling precise identification of SAMs (Figures S2A-S2D). Numerous SAMs were localized around sections of the leptomeningeal arteries with Aβ deposits, at a significantly higher density than in sections of arteries without Aβ deposition (Figure 2B). Higher magnification three-dimensional (3D) images showed these SAMs to be seemingly adherent to the leptomeningeal arteries. Furthermore, the number of SAMs adjacent to the artery wall was strongly and directly correlated with the size of the Aβ artery deposits (Figure 2C), with a significantly greater circularity index than SAMs not adjacent to Aβ deposits, suggesting a morphological transition of SAMs associated with or upon adherence to Aβ-deposited arteries (Figures 2D, 2E, S2E, and S2F).

**Figure 2.**
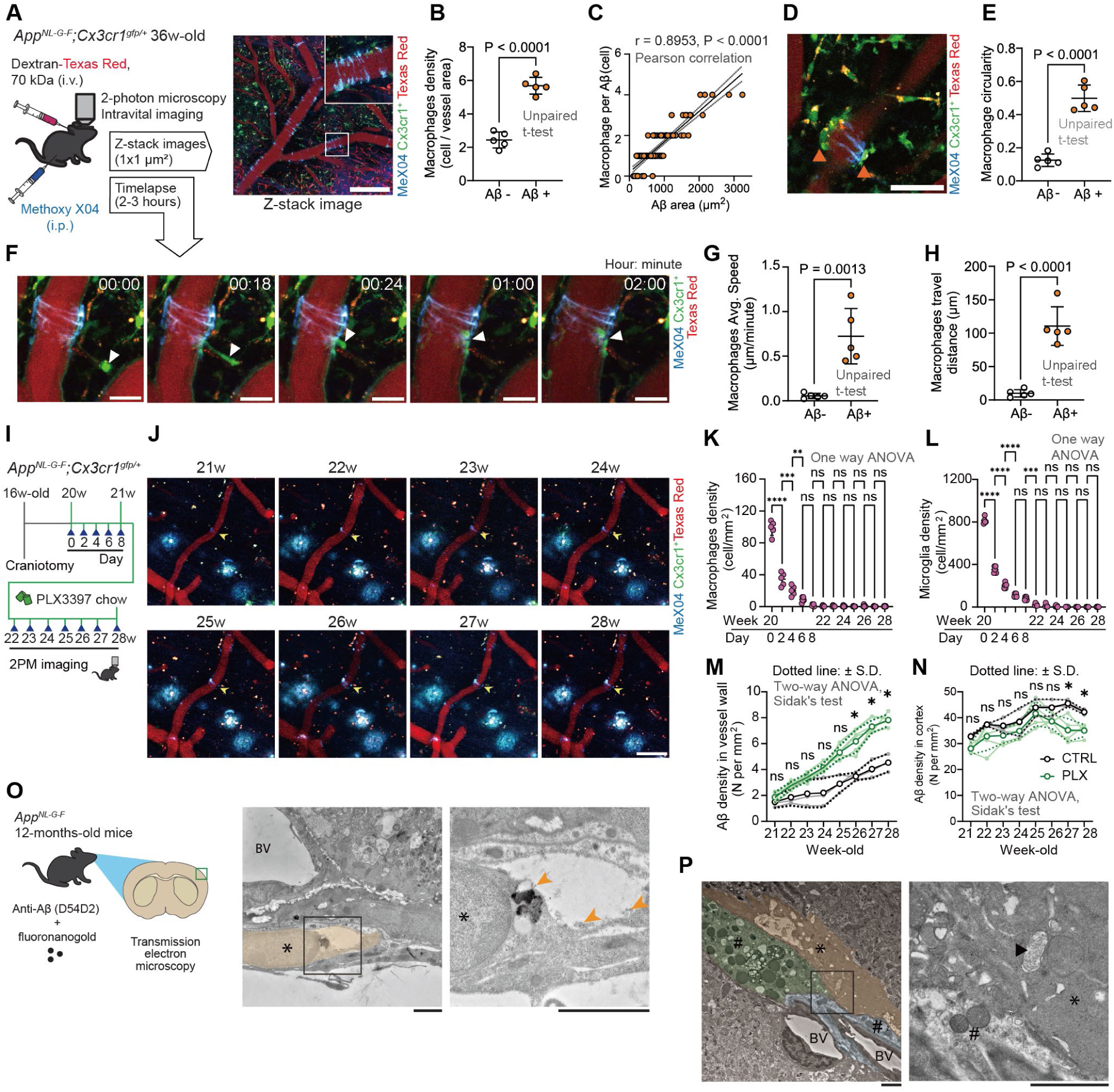
Macrophage reactivity to subarachnoid Aβ deposits. **(A)** Experimental scheme of time-lapse in vivo two-photon imaging of *App^NL-G-F/NL-G-F^ Cx3cr1^GFP/+^* after injection with Methoxy-X04 intraperitoneally and 70 kDa Texas red dextran intravenously to visualize the Aβ deposition and vessels, respectively. A typical image showing an Aβ-positive vessel with an attached macrophage. **(B)** Quantification of macrophage density between Aβ-positive and Aβ-negative vessel areas. **(C)** Correlation between the Aβ burden and the number of macrophages adhered to the Aβ-positive vessel surface. **(D and E)** Morphological comparison of macrophages that adhered to Aβ-positive and Aβ-negative vessels. **(F)** Typical time-lapse images of the immune response of subarachnoid macrophages. **(G and H)** Quantitative analysis of time-lapse imaging reveals that macrophages responding to Aβ deposition exhibited a significantly greater average speed and travel distance than did non-responsive macrophages, indicating active migration toward Aβ-positive vessel regions. **(I)** Experimental scheme of the macrophage and microglia pharmacological ablation using PLX3397. **(J)** Typical time course image of Aβ deposition, and microglia and macrophage depletion in the brain and subarachnoid space. **(K and L)** The number of macrophages and microglia was significantly reduced with PLX3397 administration. **(M and N)** Summary graph of Aβ deposition density in the subarachnoid space and brain parenchyma. **(O)** Ultrastructure of a brain macrophage (*) with an Aβ signal (pointed with orange arrow as black dot). **(P)** Electron micrograph showing a macrophage (*) located adjacent to vascular smooth muscle cells (#). A phagosome (black arrow) containing cytoplasmic material from a neighboring cell is visible within the macrophage. Data are given as mean ± standard deviation (b, e, g, h, k–n). Statistical comparisons were made using unpaired t-tests (B, E, G, H), or two-way ANOVA followed by Sidak’s test (K–N). ns = not significant, * = p < 0.05, ** = p < 0.01, *** = p < 0.001, **** = p < 0.0001. Each dot and line represents individual replicates. Scale bar, 2 µm (O, P), 25 µm (J), 50 µm (D, F), and 250 µm (A).

Time-lapse imaging directly captured SAMs migrating toward regions of Aβ deposition and adhering to the leptomeningeal artery wall (Figure 2F; Video S2). In the example shown in Figure 2F, a single SAM migrated from the subarachnoid space to the artery wall in approximately 30 min. These migrating SAMs had significantly greater velocity and travel distance than basal motility in SAMs located in unaffected vessel areas (Figures 2G and 2H). This suggests active recruitment of Aβ deposits in leptomeningeal arteries.

Subsequently, we asked whether SAMs may influence the pattern or extent of Aβ deposition. To answer this, we pharmacologically ablated macrophages through daily administration of the colony-stimulating factor 1 (CSF-1) receptor inhibitor, PLX3397. PLX3397 (600 mg/kg; MedChemExpress, NJ, USA) was added to the food (Research Diets Inc., New Brunswick, NJ, USA) of 20-week-old mice for 8 weeks (Figure 2I). The CSF-1 receptor is required for microglia survival; therefore, PLX3397 treatment almost eliminated macrophages within the subarachnoid space and cortical parenchyma within the first week and was maintained over 8 weeks (Figures 2J–2L). The number of Aβ deposits within the leptomeningeal arteries was increased in macrophage-depleted mice (Figure 2M). In contrast, the density of Aβ deposits in the cortical parenchyma modestly decreased with sustained microglial depletion (Figure 2N), as previously ascribed to the facilitation of plaque formation induced by microglial inflammation ^30^. Therefore, SAMs play a distinct role from microglia in the extent of Aβ deposition in leptomeningeal arteries. Based on the above, we hypothesized that SAMs phagocytosed arterial Aβ deposits. Subsequently, transmission electron microscopy (TEM) of cortical surface thin brain sections from *App^NL-G-F^* mice stained with an anti-Aβ antibody linked with gold particles identified macrophages adjacent to leptomeningeal arteries that contained an internalized Aβ signal within their lysosomes (Figure 2O). Notably, the macrophages displayed a distinct phagosome with engulfed cytoplasmic material, potentially derived from an adjacent vascular smooth muscle cell (Figure 2P). This finding suggests macrophage-mediated Aβ phagocytosis within the perivascular niche, including vascular smooth muscle, with potential functional effects on hemodynamics. We performed parabiosis between *App^NL-G-F^* and control mice to determine whether some of the Aβ deposits may originate from the systemic circulation. No Aβ deposition was detected in control mice, suggesting that Aβ originated from the brain parenchyma (Figures S3A-S3C).

### Transcriptome analysis of SAMs in App^NL-G-F^ mice

We undertook a phenotypic characterization of SAMs in AD mice by isolating subarachnoid meninges and adjacent parenchymal tissues for cell sorting and subsequent bulk and single-cell RNA sequencing (scRNA-seq). We carefully dissected only the thin, superficial layer of the brain surface, minimizing the deeper perivascular compartments, and performed a multi-color flow cytometry gating strategy designed to enrich for SAMs (CD11b^+^/ CD45^high^/ Tmem119^low^/ CD206^high^), and minimize the presence of parenchymal microglia (Figures 3A, 3B, S4A-S4C, S5A, and S5B). To identify early phenotypic changes, we compared the transcriptomes of isolated SAMs from 28-week-old homozygous *App^NL-G-F^* mice with age-matched control mice. Differential expression analysis of 24,395 genes revealed 299 genes that were significantly upregulated and 148 genes that were downregulated in *App^NL-G-F^* SAMs relative to controls (Figure 3C). Gene ontology enrichment analysis identified significant enrichment of genes associated with four signaling pathways (Figure 3D), including those associated with the lysosome and those associated with collagen-containing extracellular matrix (ECM) components (Figure 3E). Notably, genes encoding some of the cathepsin family of enzymes (*Ctsd, Ctsl, Ctsz),* common to both pathways, were upregulated. The pathway analysis suggests a disease-specific phenotype extending to AD SAMs, with specific collagen and lysosome signaling gene families consistent with SAM-mediated Aβ phagocytosis, smooth muscle engulfment, and extracellular tissue remodeling, which may promote the structural and hemodynamic changes associated with Aβ deposits in affected cerebral arteries (Figure 3F).

**Figure 3.**
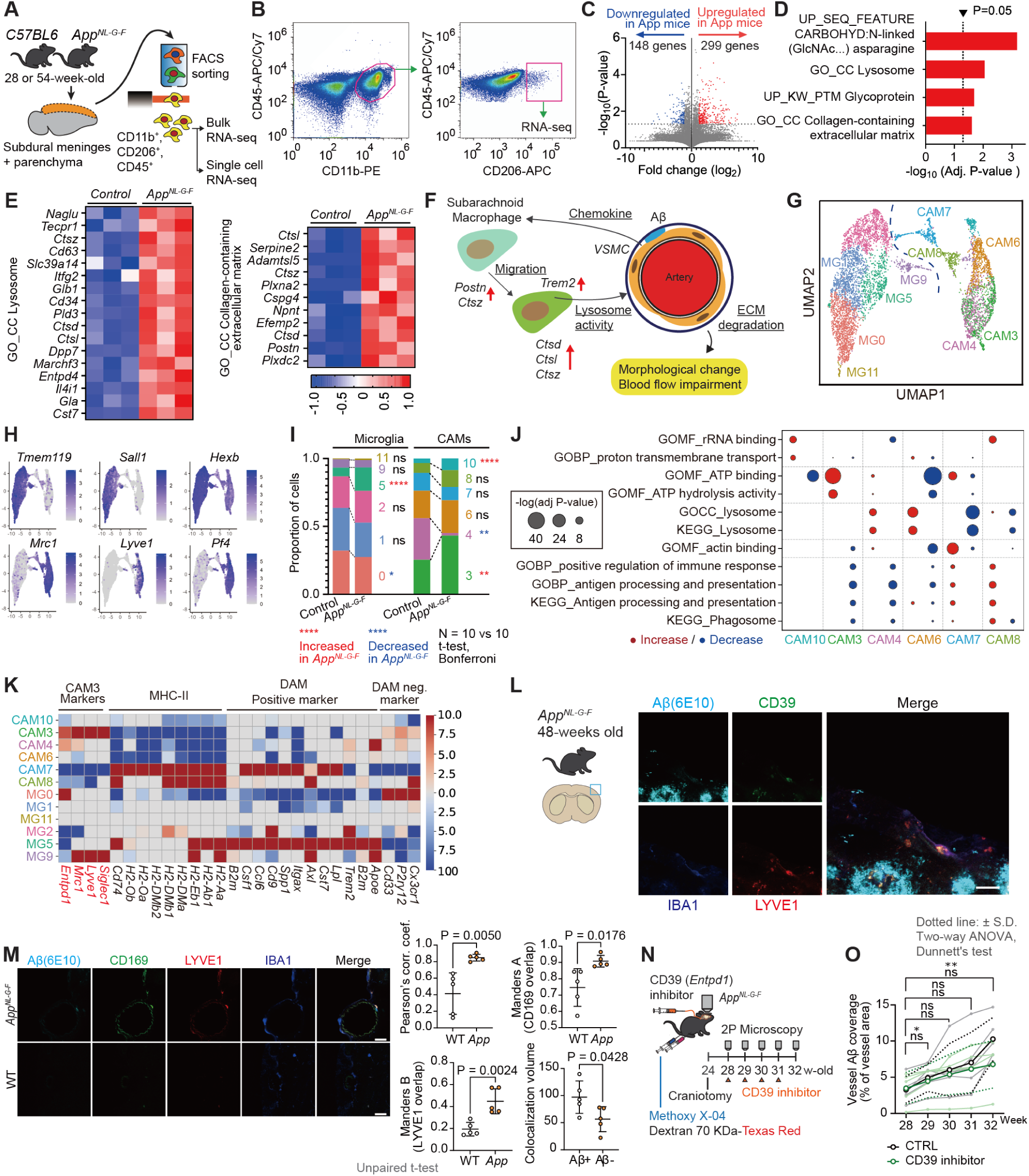
Transcriptome analysis of SAMs in *App^NL-G-F^* mice. **(A)** Experimental scheme of single-cell collection from *App^NL-G-F^* and *C57BL/6J* mice. **(B)** Gating strategy to isolate CD206^+^ macrophages from brain tissue. **(C)**, Volcano plot of RNA expression in CD206^+^ macrophages reveals upregulated and downregulated genes. **(D)** Gene sets obtained by functional pathway enrichment analysis of differentially expressed genes. (E) Heatmap of genes related to the lysosome and collagen-containing extracellular matrix. **(F)** Subarachnoid macrophages may migrate toward blood vessels with Aβ deposition, leading to extracellular matrix degradation and vascular wall remodeling, potentially impairing local cerebral blood flow. (**G**–**K**) Single-cell transcriptomic landscape of cortical myeloid cells reveals disease-associated macrophage subsets. (**G**) Uniform manifold approximation and projection showing microglia (blue) and macrophage (orange) clusters. (**H)** Expression of canonical markers (*Tmem119*, *Sall1, Hexb* for microglia; *Mrc1, Lyve1*, *Pf4* for macrophages) confirms cell-type. **(I)** Relative proportions of clusters in the *C57BL6J* and *App^NL-G-F^* cortex reveal increased clusters 3, 6, and 10 (disease-associated macrophages) and decreased cluster 0. **(J)** Functional annotation of differentially expressed genes across clusters shows that cluster 3 macrophages share immunosuppressive and ATP-metabolic features with cluster 0 microglia, whereas cluster 10 macrophages are enriched for ribosomal/mitochondrial genes. **(K)** Cluster 3 expresses *Lyve1, CD169, CD39*, and reduced MHC class II genes, consistent with an immunosuppressive phenotype. (**L, M**) Immunofluorescence confirms increased expression of Lyve1, CD169, and CD39 in *App^NL-G-F^* brains, validating disease-associated CAMs (DA-CAMs). Left top graph: Pearson’s coefficient shows the overall correlation between CD169 (488 chA) and LYVE1 (555 chB) in the IBA region. Right top graph: the fraction of CD169 signal that overlaps with LYVE1 inside the IBA1 region. Bottom left graph: The data shows the fraction of LYVE1 signal that overlaps with CD169. Bottom right graph: Colocalized surface created from the CD169–LYVE1 overlap inside IBA1, then compared this surface with the Ab surface. Ab⁺ = colocalization surface overlaps with Ab (> 0). Ab⁻ = no overlap with Ab (= 0). These cell populations increased in *App^NL-G-F^* brains and were specifically located in leptomeningeal arteries. **n**, Experimental scheme. *App^NL-G-F^* mice underwent cranial window surgery at 24 weeks of age, and a cannula was implanted for direct infusion into the cisterna magna. Mice were treated with a CD39 inhibitor (ARL67156) or vehicle control. Longitudinal two-photon imaging was performed weekly from 28 to 32 weeks of age. **o**, Quantification of vessel-associated Aβ coverage over time in control and CD39 inhibitor-treated mice. Data are presented as mean ± standard deviation (**N**). Statistical comparisons were made using two-way analysis of variance followed by Dunnett’s test (**K**–**N**). ns = not significant, * = p < 0.05 vs. control group, ** = p < 0.01 vs control group. Each dot and line represents individual replicates. Scale bar, 20 µm (**l**, **m**).

To investigate the functional heterogeneity of cortical myeloid cells interacting with Aβ, we performed scRNA-seq of microglia and CAMs isolated from the brain cortex of control and *App^NL-G-F/NL-G-F^* mice using the 10x Chromium platform (Figures S5C and S5D). After quality control, unsupervised clustering of the remaining 5941 cells combined with uniform manifold approximation and projection resulted in 12 transcriptionally distinct clusters, which constitute two major cellular compartments: microglia expressing *Tmem119*, *Sall1,* and *Hexb*, and CAMs characterized by *Mrc1*, *Lyve1*, and *Pf4* expression (Figures 3G and 3H). A comparison of the cluster composition between control and *App^NL-G-F/NL-G-F^* mice showed that several clusters were significantly altered in AD, including an increased microglial cluster (MG5; DAM-like) and three CAM clusters (clusters CAM3, CAM6, and CAM10), which we defined as disease-associated macrophage (DA-CAM) populations (Figures 3I and S3E).

Functional annotation of differentially expressed genes across clusters revealed that the DA-CAM cluster CAM3 exhibited upregulation of genes related to adenosine triphosphate (ATP) metabolism and downregulation of immune activation pathways, including those involved in Major Histocompatibility Complex (MHC) class II antigen presentation (Figure 3J). Notably, cluster CAM3 shared functional signatures with the downregulated microglial cluster MG0, suggesting a convergent immunosuppressive and metabolic phenotype between these distinct myeloid subsets (Figures S5F–S5I). In contrast, cluster CAM10 showed strong enrichment of ribosomal and mitochondrial gene sets, indicating a disease-associated activation state not observed in control mice.

Marker analysis highlighted *Lyve1, CD169, and CD39*, as well as reduced MHC class II-associated gene expression, as characteristic features of cluster 3 macrophages and the analogous microglial subset. The proximity of our DAM-like microglia cluster (cluster MG5) to previously reported DAM signatures confirmed the robustness of our analysis pipeline (Figure 3K). Immunofluorescence staining validated the increased expression of Lyve1, CD169, and CD39 in *App^NL-G-F/NL-G-F^* mouse brains, confirming the presence of a DA-CAM characterized by ATP catabolism and immunosuppressive function (Figures 3L and 3M). Notably, these cells were specifically increased and localized adjacent to the leptomeningeal arteries in *App^NL-G-F/NL-G-F^* mice (Figure 3M).

Disease-associated molecular programs or phenotypes in immune cells are often associated with ATP, which generally stimulates a pro-inflammatory transition. Conversely, adenosine maintains a basal or anti-inflammatory state. A range of ecto-nucleotidases on the surface of myeloid cells, including CD39 (encoded by *Entpd1*), contribute to local ATP metabolism, thereby acting as “immune suppressors” in regulating immune cell polarization. We hypothesized that inhibiting this immunosuppressor may enhance the capacity of SAMs to phagocytose Aβ in the subarachnoid space, thereby reducing arterial deposits. We targeted CD39 because an effective pharmacological inhibitor (ARL67156) ^31^ was available, and various CD39 inhibition strategies are currently in clinical trials for cancer. ARL67156 was administered intracisternally ^32^ to ensure exposure to SAMs, as this nucleotide analogue has limited BBB permeability. ARL67156 was injected directly into the CSF via an indwelling catheter in the cisterna magna, with weekly injections over 4 weeks combined with two-photon imaging to track Aβ artery deposition (Figure 3N). We focused on 28–32-week-old mice, when deposits were just becoming apparent, to avoid excessive Aβ deposits, smooth muscle loss, and vascular complications. Successful delivery was confirmed using fluorescein-conjugated dextran (Figure S5J). We found that vascular Aβ coverage significantly increased after week 28 in control mice but remained unchanged with CD39 inhibition (Figure 3O).

### Hemodynamic consequences of Aβ deposition and macrophage polarization

Given our preceding findings, we hypothesized that these immune–vascular interactions compromise arterial contractility and cerebral perfusion. To correlate Aβ deposition, SAM migration, and polarization to hemodynamic parameters, we initially used *in vivo* imaging to quantify leptomeningeal vascular parameters (Figure 4A). Vessel diameter was consistently larger in segments of arteries with Aβ deposits than in regions without (Figures 4B-4D). Time-lapse imaging (15 frames/s) was performed to observe the dynamic changes in the diameter of Aβ-associated artery segments (Figure 4E). Diameter fluctuations arise from various sources with different frequencies; in particular, a Fast Fourier transform spectra of diameter changes within the range we could resolve (≤7.5 Hz) identified a marked reduction in the amplitude of slow fluctuations (≤0.5 Hz, Figure 4F), which is important for perivascular clearance flow ^33,34^. Reduced diameter fluctuations in Aβ-associated arteries were, however, observed across the entire frequency range, suggesting a loss of compliance and/or vasomotor contractility.

**Figure 4.**
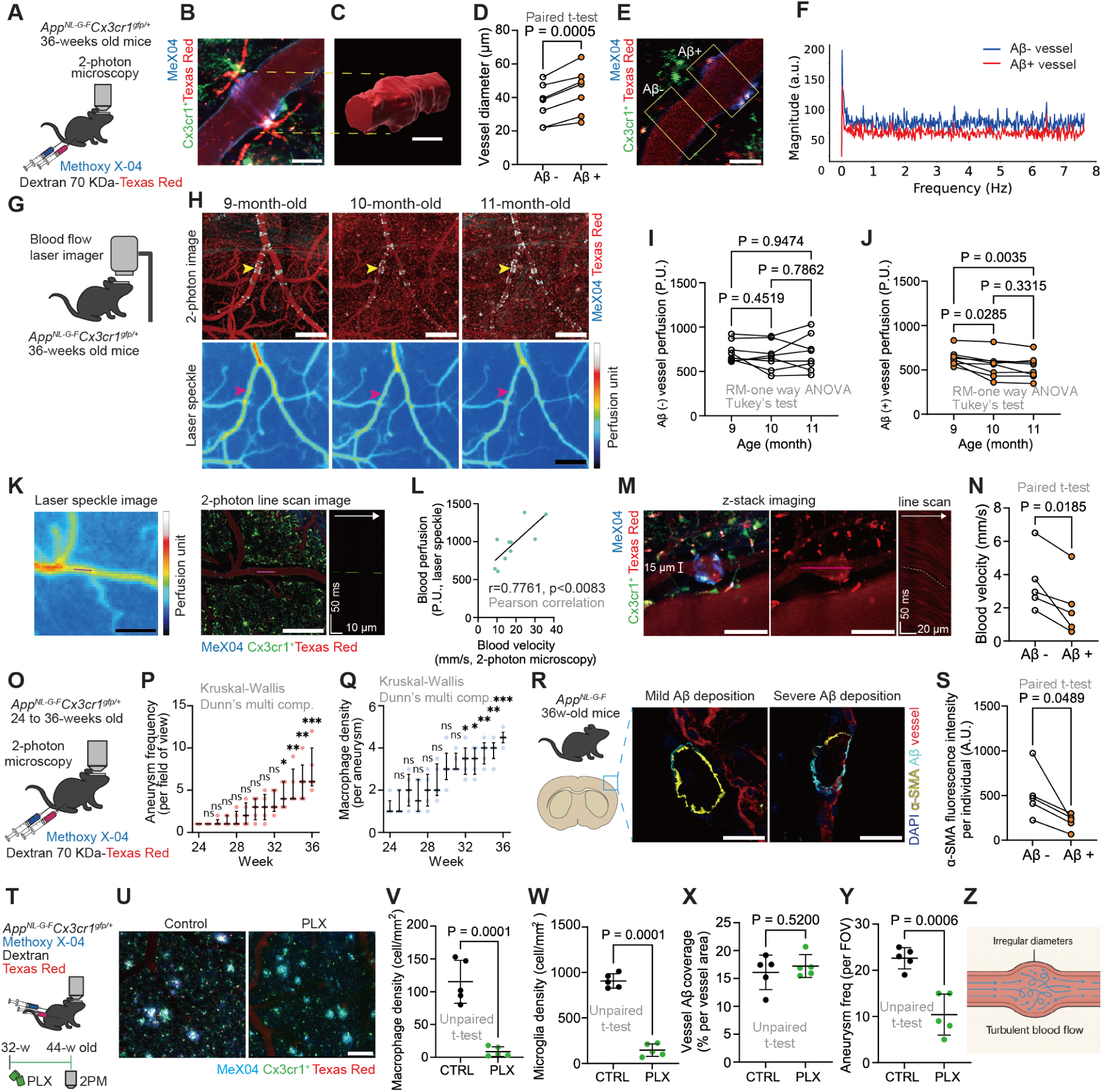
Hemodynamic consequences of Aβ deposition. (**A**) Experimental scheme of the two-photon imaging to measure the vessel diameter and its changes. **(B)** Typical *in vivo* image of a blood vessel with Aβ deposition in *App^NL-G-F/NL-G-F^Cx3cr1^GFP/+^* mice. **(C)** 3D reconstruction of the image in B. **(D)** Summarized graph of the vessel diameter. **(E)** Region of interest: Aβ deposition-positive vessel region and Aβ deposition-negative region were used to measure their contraction. **(F)** Fast Fourier transform spectra of vessel diameter changes show lower signal magnitude in Aβ-positive (red) than in Aβ-negative (blue) vessels. **(G)** Experimental scheme of laser speckle imaging to measure cerebral blood perfusion. **(H)** Typical two-photon images showing an Aβ-positive vessel and respective laser speckle images at different time points. (**I, J**) Summarized graph shows cerebral vessel blood perfusion. **(K)** Representative images of laser speckle and two-photon line scans. (**L)** Quantification of laser speckle and two-photon line scan correlation data. **(M)** Typical image of a small artery with Aβ deposition, as seen in line scan imaging, shows a marked decrease in blood flow. **(N)** Quantification of blood flow velocity using line scan imaging. **(O)** Experimental scheme for aneurysm observation in *App^NL-G-F/NL-G-F^Cx3cr1^GFP/+^* mice. (**P**, **Q)** Summarized graph of aneurysm frequency and macrophage density from mice at 24 to 36 weeks of age. **(R)** Immunostaining images show cerebral vascular smooth muscle of *App^NL-G-F^* mice with mild and severe Aβ deposition. **(S)** Quantification of vascular smooth muscle in Aβ-positive and Aβ-negative vessels. **(T)** Experimental design. *App^NL-G-F/NL-G-F^Cx3cr1^GFP/+^* mice were fed with standard feed or PLX3397-containing chow for 12 weeks, starting at 32 weeks of age. **(U)** Representative *in vivo* two-photon images at 44 weeks of age showing leptomeningeal vessels (magenta), Aβ deposits (cyan), and Cx3cr1⁺ cells (green) in control and PLX-treated mice. (**V**–**Y**) Quantification of subarachnoid macrophage density, cortical microglia density, vessel Aβ coverage, and aneurysm frequency per field of view. **(Z)** Illustration of turbulent flow in a vessel with an irregular diameter. Statistical analysis was performed using paired t-tests (**D**, **N**, **S**) and repeated-measure one-way analysis of variance followed by Tukey’s test (I, J), Kruskal–Wallis test (**P**, **Q**), and unpaired t-tests (**V**–**Y**). Error bars indicate the mean ± standard deviation; ns = not significant, *p < 0.05, **p < 0.01, ***p < 0.001. Scale bar, 25 µm (**B**, **C**, **E**, **P**), 50 µm (**L**), 100 µm (**U**), 200 µm (**K**), 250 µm (**H**).

To better determine how vascular changes affect tissue perfusion, we employed two approaches based on red blood cell flow in arterioles (Figure 4G). Laser speckle contrast imaging (LSCI) uses proprietary software to calculate arbitrary perfusion units based on differences in contrast reflection patterns within specific regions of interest. Simultaneously, two-photon line-scanning directly tracks the displacement of red blood cell shadows against the fluorescent dextran background in imaged vessel regions ^35,36^. Arteries with and without Aβ deposits were first identified at 9 months. The development of Aβ deposits was tracked over the subsequent 2 months using two-photon imaging (Figure 4H, upper panels). Blood perfusion in the same arteries was subsequently measured with LSCI (Figure 4H, lower panels). Blood flow through arteries associated with Aβ deposits significantly decreased as Aβ deposits increased, while flow in arteries not associated with Aβ deposits remained relatively stable (Figures 4I and 4J). The decline in blood flow was apparent in the first month and then stabilized, possibly reflecting chronic arterial vasodilation that compensates for reduced vasomotor activity or changes in perfusion pressure. Furthermore, LSCI was performed alongside (in separate mice) two-photon microscopy line-scanning, and as expected, there was a tight linear correlation between these two measures in a sample of larger leptomeningeal arteries (Figures 4K and 4L). Finally, using two-photon line-scanning, we confirmed that blood flow was reduced in segments of arterioles with Aβ deposits than in segments in the same arterioles without Aβ deposits (Figures 4M and 4N).

The results suggest a sequence of vasomotor deficits associated with Aβ deposits, macrophage recruitment, and arterial dilation, followed by subsequent decreases in cerebral perfusion as vascular Aβ deposits and inflammation progress. Leptomeningeal arterial aneurysms were consistently increased during imaging, paralleled by an increased density of macrophages around the aneurysm sites (Figures 4M and 4O-4Q). Given the presence of putative smooth muscle fragments within macrophage phagosomes, we hypothesized that aneurysms and other vascular abnormalities are related to macrophage-mediated loss of smooth muscle due to Aβ deposits. Consistently, we observed a marked loss of smooth muscle actin immunostaining in arteries with Aβ deposition compared with those without in 36-week-old mice (Figures 4R and 4S). This strongly suggests that vascular and blood flow deficits arise due to smooth muscle cell loss.

We further examined whether depletion of microglia and SAMs influences the formation of vascular aneurysms. We administered PLX3397 chow to 32-week-old *App^NL-G-F^; Cx3cr1^gfp/+^* mice. Using two-photon *in vivo* imaging, we found that PLX3397 administration effectively depleted microglia and SAMs (Figures 4T-4W). Vessel Aβ coverage remained unchanged, likely because amyloid deposition at 44 weeks of age had reached a saturated stage of progression (Figure 4X). Notably, PLX3397-treated mice showed a reduction in the frequency of vascular aneurysms compared with controls (Figure 4Y). Therefore, the differential effects of SAMs include facilitating the formation of vascular aneurysms around Aβ-laden arteries and reducing Aβ deposition at earlier stages. This dual effect may explain why pharmacological depletion of macrophages had a minimal impact on blood flow and brain necrosis (Figures S6A-S6D and S7A-S7C), suggesting that more targeted therapy is needed. These findings indicate that immune cell-mediated mechanisms and Aβ accumulation, may contribute to the structural vulnerability of leptomeningeal vessels and impair blood flow (Figure 4Z; summarized scheme in Figure S8).

## DISCUSSION

AD is characterized by the progressive accumulation of Aβ plaques and neurofibrillary tangles, resulting in a progressive cognitive decline and dementia.^19,37^ A common comorbidity of AD is the development of Aβ plaques around cerebral vessels (CAA), which initially appear in the pial and subarachnoid arteries and subsequently spread to the parenchymal arteries.^38^ As the extent of deposits further develops, impaired perfusion and hemorrhages are detected throughout the parenchymal and leptomeningeal arteries, contributing to cognitive and neurological sequelae. The central hallmark of these diseases is reduced blood flow in the brain. The loss of arterial smooth muscle is associated with vascular deficits in advanced CAA and AD. However, the mechanism by which Aβ deposits cause these cellular and functional deficits remains unclear. Therefore, we undertook time-lapse imaging to track the formation and development of Aβ deposits in subarachnoid vessels and their functional consequences, with a particular focus on interactions with the surrounding immune environment.

Our time-lapse imaging and 3D reconstruction demonstrate banded Ab deposits that form circumferential rings or coils around the leptomeningeal arteries. These leptomeningeal arteries are highly vulnerable to Ab, with vessel deposits even preceding the development of significant Ab plaques in the underlying parenchymal cortex. At arterial wall locations where these Ab coils form, there is a loss of smooth muscle actin, and macrophages become embedded within the arterial wall. Here, we identify the nature of these macrophage interactions. Within the subarachnoid space reside a discrete population of CAMs that adhere to vessels, trabeculae, and meninges, defined as SAMs.^39^ We directly observed these SAMs actively migrating to and accumulating around the Ab deposits, undergoing concurrent polarization associated with a more activated and phagocytic phenotype. Remarkably, electron microscopy revealed both Aβ clusters and smooth muscle cell components within the phagocytic lumen of these macrophages, and depleting the macrophage population increased the formation of vessel Aβ deposits. This provides strong evidence that SAMs are directly involved in the regulation and clearance of leptomeningeal cerebral Ab deposits.

Recent transcriptomic and fate-mapping studies have demonstrated that CAMs are distinct from peripheral macrophages and microglia. Information on how these transcriptomic brain CAM clusters transform during disease states is limited. Our bulk tissue RNA analysis revealed upregulation of lysosomal and ECM pathways, consistent with our observations of macrophage migration, adherence to the arterial vessel wall, and phagocytosis. However, this bulk tissue analysis was unable to identify cell-specific signaling pathways that may trigger this response. However, the scRNA-seq clearly distinguished microglial and macrophage populations within this tissue sample and identified a specific macrophage cluster (CAM3) that characterized the SAMs surrounding arterial Ab deposits. CAM3 showed increased ATP metabolism and signaling pathways associated with disease or damage phenotypes and was significantly upregulated in the mouse model of AD. This cluster likely corresponds to the previously identified dominant MHC-low SAM cluster,^1,16^ which, like parenchymal microglia, is self-renewing and resides in the brain from the early embryonic period. These SAMs appear to expand in AD, as previously reported for microglia DAM clusters^5^ (our MG5 cluster). In addition, these SAMs express CD39 (*Entpd1*), and perfusion of the CD39 inhibitor into the CSF reduced the development of arterial Ab deposits. Therefore, we identified a specific SAM-immune suppressor molecule that could be targeted for therapeutic intervention in CAA.

Ultimately, the cognitive decline from CAA and AD-associated vascular Aβ deposits arises from reduced cerebral tissue perfusion owing to parenchymal hemorrhages associated with more severe AD and CAA,^40,41^ resulting from a severe loss of smooth muscle in parenchymal arterioles and loss of arterial compliance and/or vasomotor reactivity. We present an earlier sequence of events in subarachnoid arterioles that may precede these parenchymal events. Aneurysms frequently formed in leptomeningeal arteries at sites of SAM accumulation, and we directly observed reduced red blood cell flow through these vessel protrusions. Turbulent blood flow may reduce the perfusion pressure supplied to the penetrating parenchymal arterioles. Vasodilation, or other physiological adaptations, may compensate for the reduced perfusion pressure; however, as the smooth muscle of parenchymal arterioles and vasomotor capacity progressively diminish, this compensation fails, and cerebral perfusion drops. Depleting SAMs reduced these leptomeningeal aneurysms, suggesting that SAMs contribute to the elimination of arterial Aβ deposits.

We have identified a discrete population of SAMs as key mediators of vascular defects in leptomeningeal arteries with early Ab deposits. While their restricted location within the subarachnoid space may enhance selective pharmacological targeting, leveraging these SAM signaling pathways to mitigate against vascular and neurological sequelae in AD and CAA will need to be nuanced. The incidence of ARIAs associated with broad Aβ antibody targeting underscores the need for thoughtful approaches that account for vascular complications. Enhanced SAM phagocytosis of Aβ deposits, for example, via a CD39 inhibitor, may also result in the loss of vessel smooth muscle and the development of localized weakened artery regions prone to aneurysms. Careful consideration of timing may be needed, with potentially early CD39 inhibition before extensive deposit formation. In addition, CD39 inhibitors may find a place alongside Aβ antibody therapy to help clear brain Aβ while minimizing the resultant potential vascular complications.

## RESOURCE AVAILABILITY

### Lead contact

Requests for further information and resources should be directed to and will be fulfilled by the lead contact, Hiroaki Wake.

### Materials availability

This study did not generate new unique reagents.

### Data and code availability

- All processed expression data supporting the findings of this study are provided in the accompanying Source Data File. All raw sequencing data (FASTQ) generated in this study have been deposited in a public nucleotide sequence repository (DNA Data Bank of Japan; a member of the International Nucleotide Sequence Database Collaboration) and will be made publicly available upon publication.
- This paper does not report original code.
- Any additional information required to reanalyze the data reported in this paper is available from the lead contact upon request.

## ACKNOWLEDGMENTS

We thank Dr Ikuko Takeda (Nagoya University) for assistance with fluorescence-activated cell sorting, and Dr Shouta Sugio (Nagoya University) for assistance with Laser Speckle Imaging. This work was supported by Grant-in-Aid for Scientific Research (A) Grant Number JP25H01217, JST, CREST Grant Number JPMJCR22P6, Japan Agency for Medical Research and Development, CREST Grant Number JP23gm1410011, programs for Bridging the gap between R&D and the IDeal society (society 5.0) and Generating Economic and social value (BRIDGE), Moonshot R&D Program Grant Number JP24zf0127012, Moonshot R&D Program Grant Number JP24zf0127010, Moonshot R&D Program Grant Number JPMJMS2012-3-4-1, Adopting Sustainable Partnerships for Innovative Research Ecosystem Grant Number JP23jf0126004, and Research Foundation for Opto-Science and Technology.

## AUTHOR CONTRIBUTIONS

Conceptualization, R.Y.H., T.T., and H.W.; methodology, R.Y.H., T.T., and H.W.; Investigation, R.Y.H., L.H., A.Y., Y.S., R.S., M.T., A.K., and T.K.; writing—original draft, R.Y.H., T.T., H.W., A.J.M., and D.L.C; writing—review & editing, R.Y.H., T.T., H.W., A.J.M., M.P., and D.L.C; funding acquisition, H.W; resources, H.W; supervision, H.W. and T.M.

## DECLARATION OF GENERATIVE AI AND AI-ASSISTED TECHNOLOGIES

During the preparation of this work, the author(s) used ChatGPT in order to generate the schematic figures. After using this tool or service, the author(s) reviewed and edited the content as needed and take(s) full responsibility for the content of the publication.

### SUPPLEMENTAL INFORMATION

**Document S1.**

**Video S1. Time-lapse imaging of ring-shaped formation of Aβ deposition around the leptomeningeal arteries, related to Figure 1M**.

Long-term imaging of the ring-shaped formation of Aβ deposition around the leptomeningeal arteries. The numbers in the top-right corner represent weeks. Scale, 25 µm.

**Video S2. Immune reaction of SAMs against Aβ deposition, related to Figure 2F**.

Immune reaction between macrophages (green) and Aβ (blue) on leptomeningeal arteries (red) was tracked in vivo over time. The numbers in the top right represent hours and minutes. Scale, 50 µm.

---

**Figure S1.**
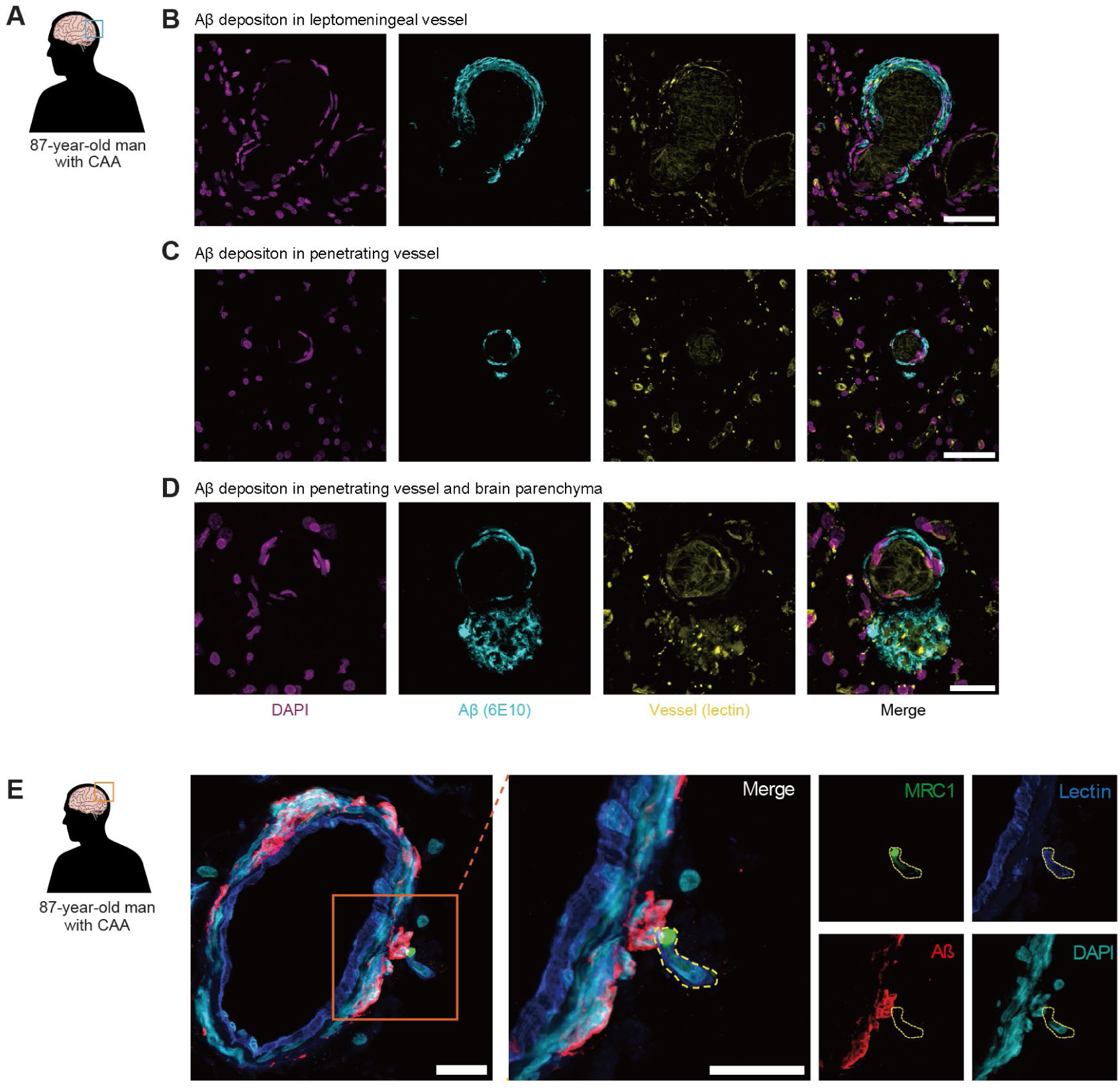
Amyloid-β (Aβ) deposition in the leptomeningeal and parenchymal compartments of a cleared human brain with cerebral amyloid angiopathy (CAA). (**A**) Immunofluorescence imaging of optically cleared post-mortem human brain tissue from a subject with CAA. (**B**) Prominent Aβ accumulation within the walls of leptomeningeal arteries. (**C**) Dense circumferential Aβ deposition around penetrating cortical arteries. (**D**) Aβ plaques in the cortical parenchyma exhibit a dense core surrounded by a diffuse region. (**E**) Representative confocal image showing a leptomeningeal vessel containing Aβ deposited along the vascular wall. A macrophage expressing MRC1 (CD206) is observed adjacent to the Aβ accumulation. Scale bar, 50 µm (B, C), 25 µm (D, E).

**Figure S2.**
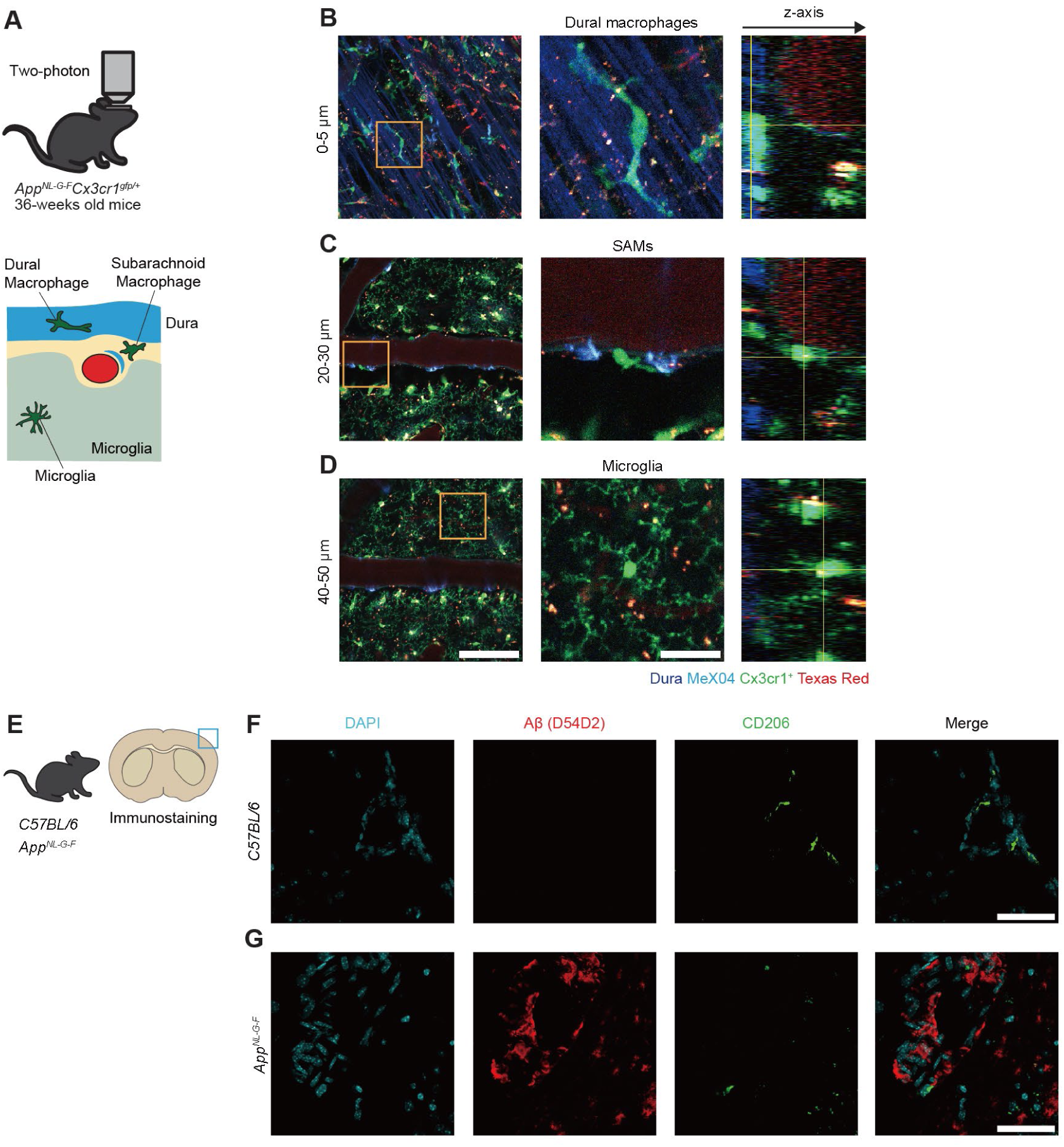
Layer-specific myeloid cell populations in the meninges and superficial cortex of *App* KI mice. (**A**) Schematic representation of CNS surface myeloid compartments: dural macrophages (top), subarachnoid macrophages (SAMs; middle), and microglia (bottom). (**B**) Maximum intensity projection of two-photon Z-stack showing myeloid cells and vasculature. Boxed regions correspond to the next panels. Dural macrophages are located at the outer brain interface and co-localized with the meningeal dura. (**C**) SAMs are located within the subarachnoid space between the dura and pia mater, with a perivascular association. (**D**) Microglia in the cortical layer exhibit a highly ramified morphology. (**E**) Immunofluorescence images of a brain surface coronal section from *C57BL/6* (wild-type) and *App^NL-G-F^* mice. (**F**) In wild-type mice, CD206^+^ macrophages are observed along surface vessels with no detectable Aβ signal. (**G**) In *App^NL-G-F^* mice, prominent Aβ deposition is detected around the brain surface vasculature and parenchyma. Scale bar, 25 µm (inset: B–D), 50 µm (F), 100 µm (B–D).

**Figure S3.**
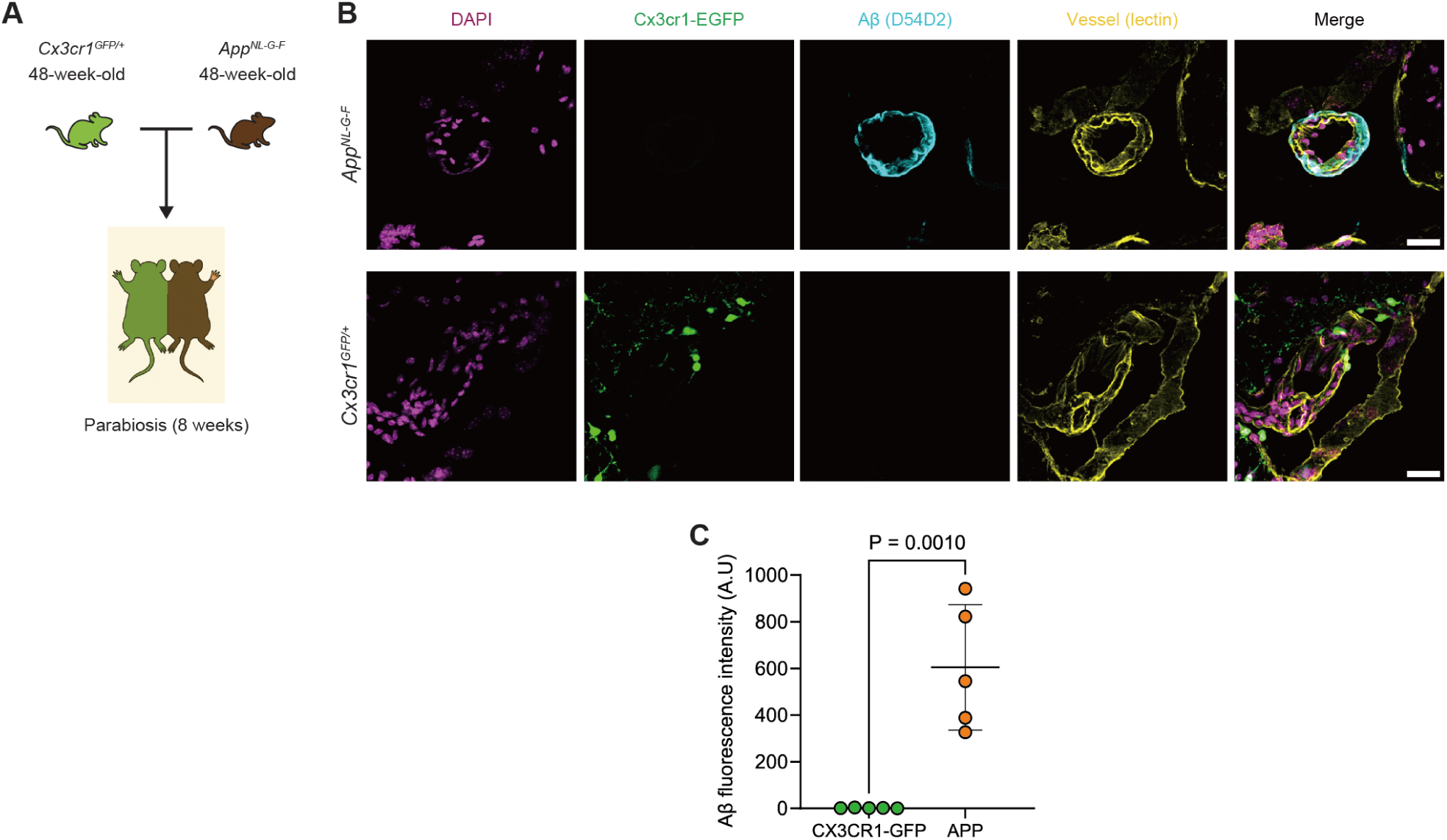
Parabiosis experiment between *App^NL-G-F/NL-G-F^* and *Cx3cr1^GFP/+^* mice. (**A**) Schematic of the parabiosis experimental design: *App^NL-G-F/NL-G-F^* mice (12 months old) were surgically joined with age-matched *Cx3cr1^GFP/+^* mice and maintained for 8 weeks. (**B**) Representative immunostaining image showing ring-shaped Aβ deposition surrounding cerebral blood vessels in *App^NL-G-F/NL-G-F^* mice, whereas no vascular Aβ was detected in the parabiotic *Cx3cr1^GFP/+^* partners. (**C**) Quantification graph summarizing the absence of Aβ accumulation in cerebral blood vessels of *Cx3cr1^GFP/+^* mice after 8 weeks of parabiosis. Data are expressed as mean ± standard deviation. Statistical comparisons were made using unpaired t-tests. N = 5 per group. Scale bar: 25 µm (B).

**Figure S4.**
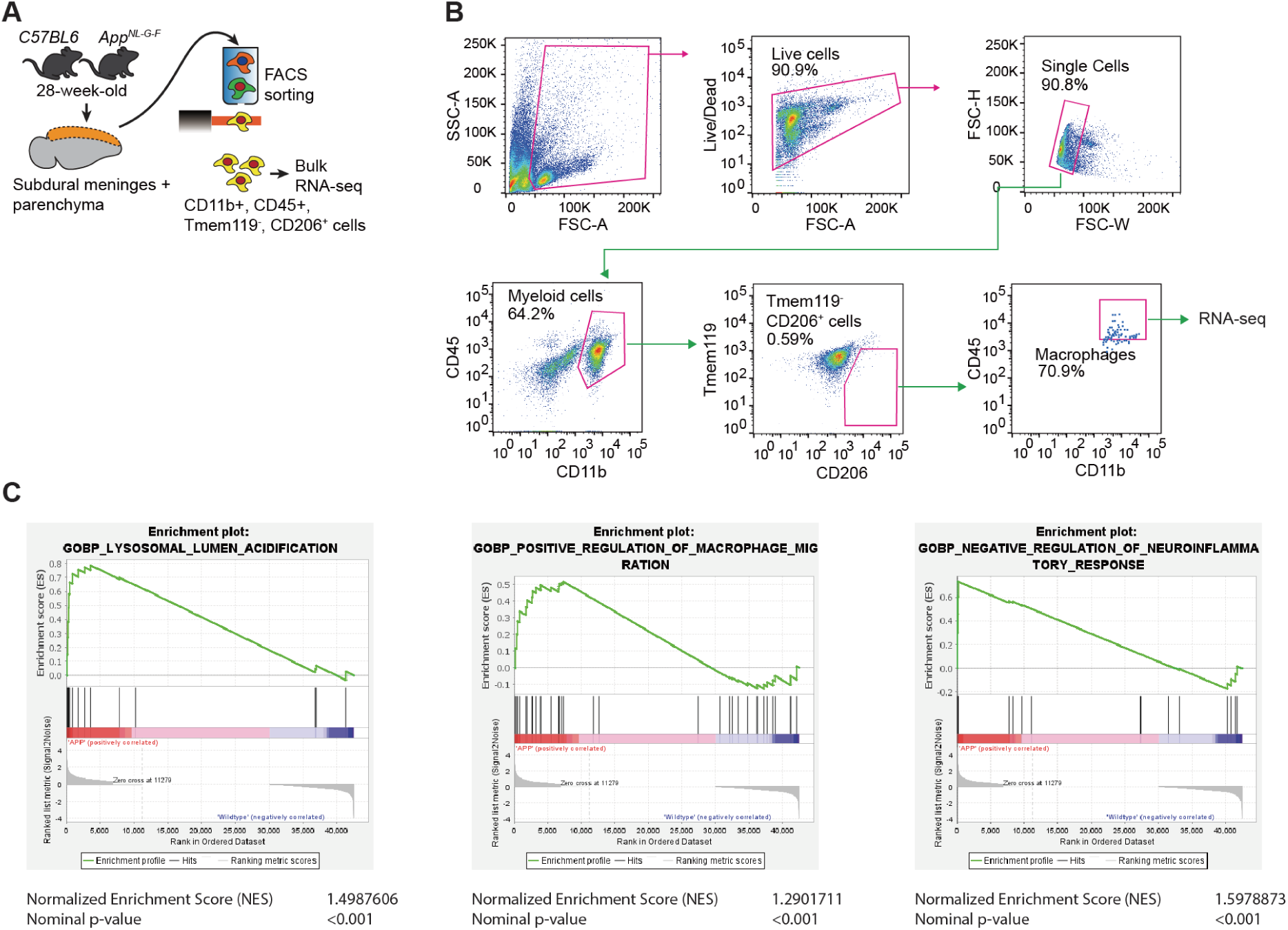
Characterization of CD206^+^ brain-associated macrophages and associated gene expression signatures. **(A)** Gating strategy for flow cytometric analysis of CD206^+^ brain-associated macrophage populations isolated from the brain parenchyma and subdural meninges of 28-week-old *App^NL-G-F^* and *C57BL/6J* mice. **(B)** CD206^+^ macrophages were defined as CD45^high^ CD11b^+^ Tmem119^−^ CD206^+^. (**C**) Gene Set Enrichment Analysis (GSEA) of sorted CD206^+^ macrophages reveals enrichment of gene sets related to lysosomal acidification (GO: Lysosomal lumen acidification), positive regulation of macrophage migration (GO: Positive regulation of macrophage migration), and negative regulation of neuroinflammatory processes (GO: Negative regulation of neuroinflammatory response). NES and p-values shown for selected gene sets.

**Figure S5.**
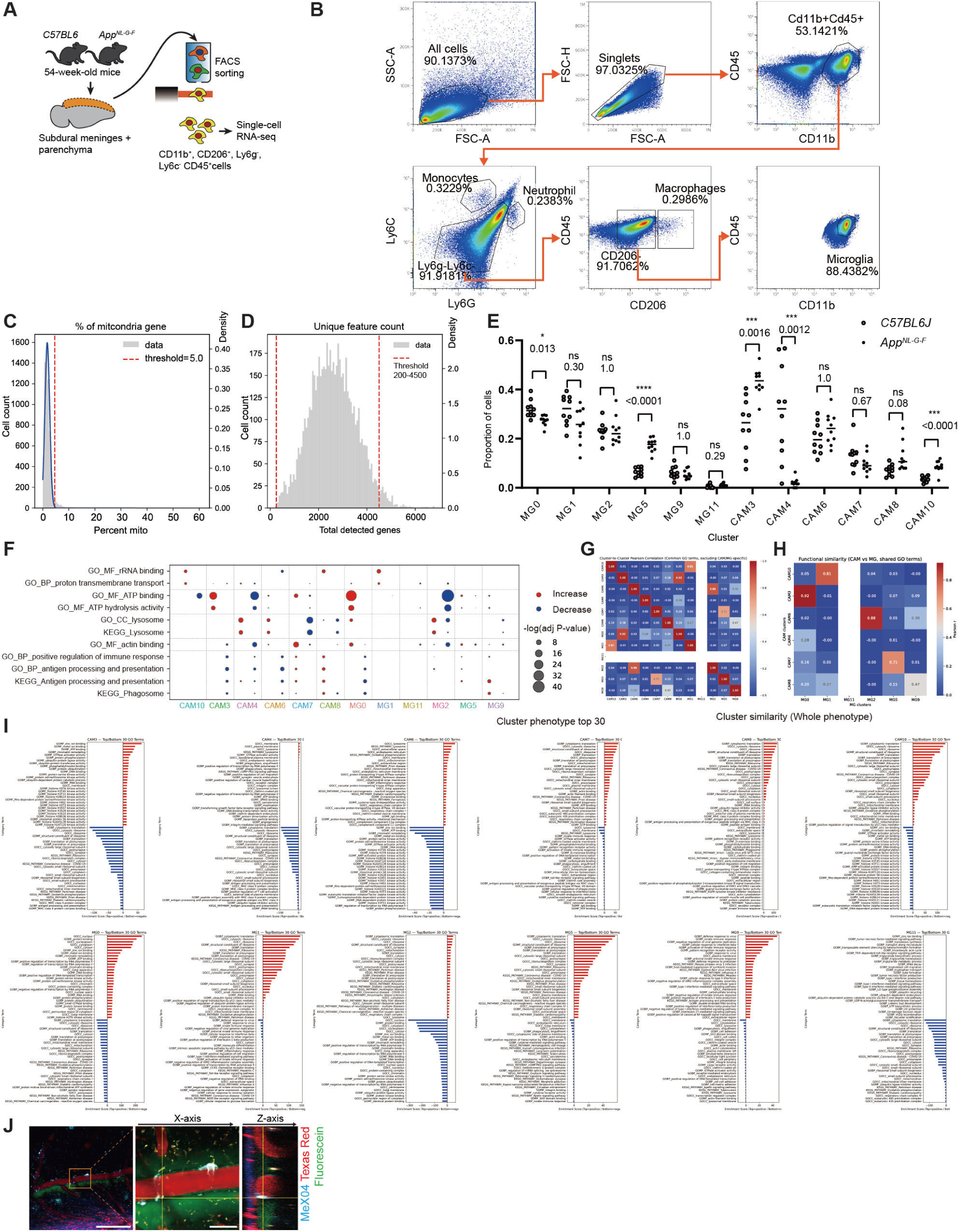
Single-cell RNA-seq supplementary analysis. (**A**) Experimental scheme of single-cell collection from *App^NL-G-F^* and *C57BL/6J* mice. (**B**) Gating strategy for flow cytometric analysis of CD206^+^ brain-associated macrophage populations isolated from the brain parenchyma and subdural meninges of 54-week-old *App^NL-G-F^* and C57BL/6 mice. (**C**) CD206^+^ macrophages were defined as CD45^high^ CD11b^+^ LY6C^−^ LY6G^−^ CD206^+^. (**D**) Cells with 200–4500 unique detected features were retained as high-quality single cells for downstream analyses. (**E**) Frequency of each microglial (MG) and CNS-associated macrophage (CAM) cluster per individual mouse. Both *C57BL/6J* and *App^NL-G-F^* groups contained *n* = 10 animals. (**F**) Functional annotation of each cell cluster was performed using DAVID based on genes significantly upregulated or downregulated in each cluster. (**G**) Correlation matrix of functional annotations showing the similarity among clusters. Only functional categories that characterized cell states beyond the MG and CAM groups were included. Pearson’s *r* values are shown. (**H**) Pairwise comparison between CAM and MG clusters. Correlated functional annotations were observed between MG0–CAM3, MG1–CAM10, MG2–CAM6, and MG5–CAM7. (**I**) Top 30 gene sets annotated by DAVID for each cluster are displayed. (**J**) Representative two-photon images showing cerebral blood vessels (Dextran–Texas Red, magenta) and Aβ deposits (Methoxy X-04, cyan), and the subarachnoid space (Dextran–Fluorescein, green).

**Figure S6.**
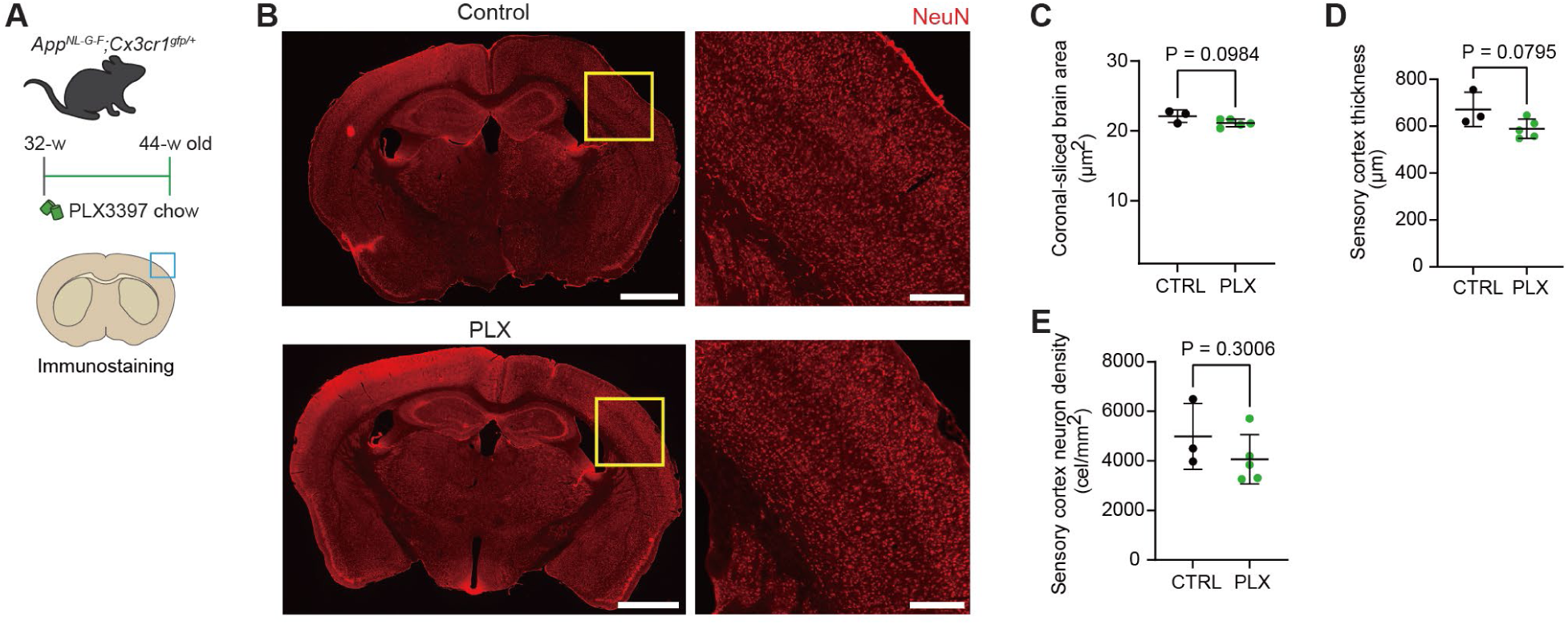
PLX treatment shows a trend toward brain shrinkage without significant neuronal loss in the sensory cortex. (**A**) Experimental design. AppNL-G-F; Cx3cr1gfp/- mice were treated with PLX3397 or a control diet for 4 weeks before perfusion and brain collection for NeuN immunostaining. (B) Representative coronal sections of the mouse brain stained with NeuN showing the neuronal distribution in control and PLX-treated groups. Yellow boxes indicate the sensory cortex region analyzed. (C) Coronal-sliced brain area in PLX-treated mice compared to that in controls (p = 0.094). (D) Sensory cortex thickness showed a non-significant trend toward reduction in PLX-treated mice (p = 0.0795). (E) Quantification of the sensory cortex neuronal density (NeuN⁺ cells/µm²) shows no significant difference between the PLX and control groups (p = 0.3006). Data are presented as the mean ± standard deviation. Statistical comparisons were made using unpaired t-tests. N = 5 per group. Scale bars, 1 mm (left panels), 200 µm (inset / right panels).

**Figure S7.**
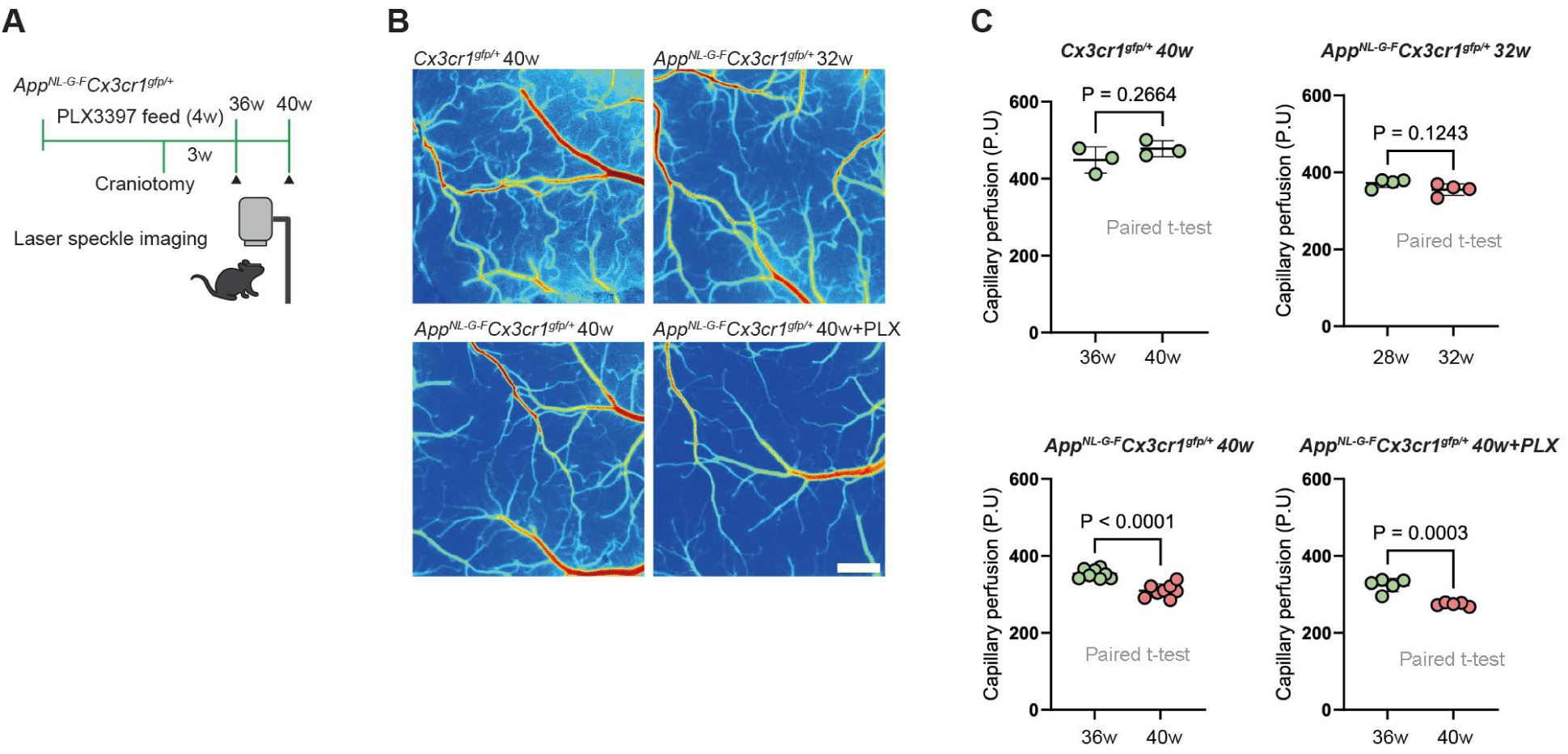
Depletion of myeloid cells by CSF1R inhibition exacerbates parenchymal capillary hypoperfusion in aged *App* KI mice. (**A**) Schematic of the experimental design showing CSF1R inhibitor (PLX3397) administration and laser speckle contrast imaging of capillary blood perfusion in *App^NL-G-F^* and *C57BL/6J* control mice. (**B**) Representative laser speckle imaging of cerebral capillary blood flow (area outside large vessels). In *C57BL/6J* and 8-month-old *App^NL-G-F^* mice, the brain parenchyma appears well perfused. In contrast, *App^NL-G-F^* mice treated with PLX3397 show marked hypoperfusion in the cortical parenchyma. (**C**) Quantification of perfused capillaries reveals no significant reduction in *C57BL/6J* mice or in young (7–8 months old) *App^NL-G-F^* mice. Meanwhile, 9–10-month-old *App^NL-G-F^* mice exhibit significant hypoperfusion, which is further exacerbated by PLX3397 treatment at the same age. Data are presented as mean ± standard deviation. Statistical comparisons were made using paired t-tests. N = 5 per group. Scale bar, 500 µm (B).

**Figure S8.**
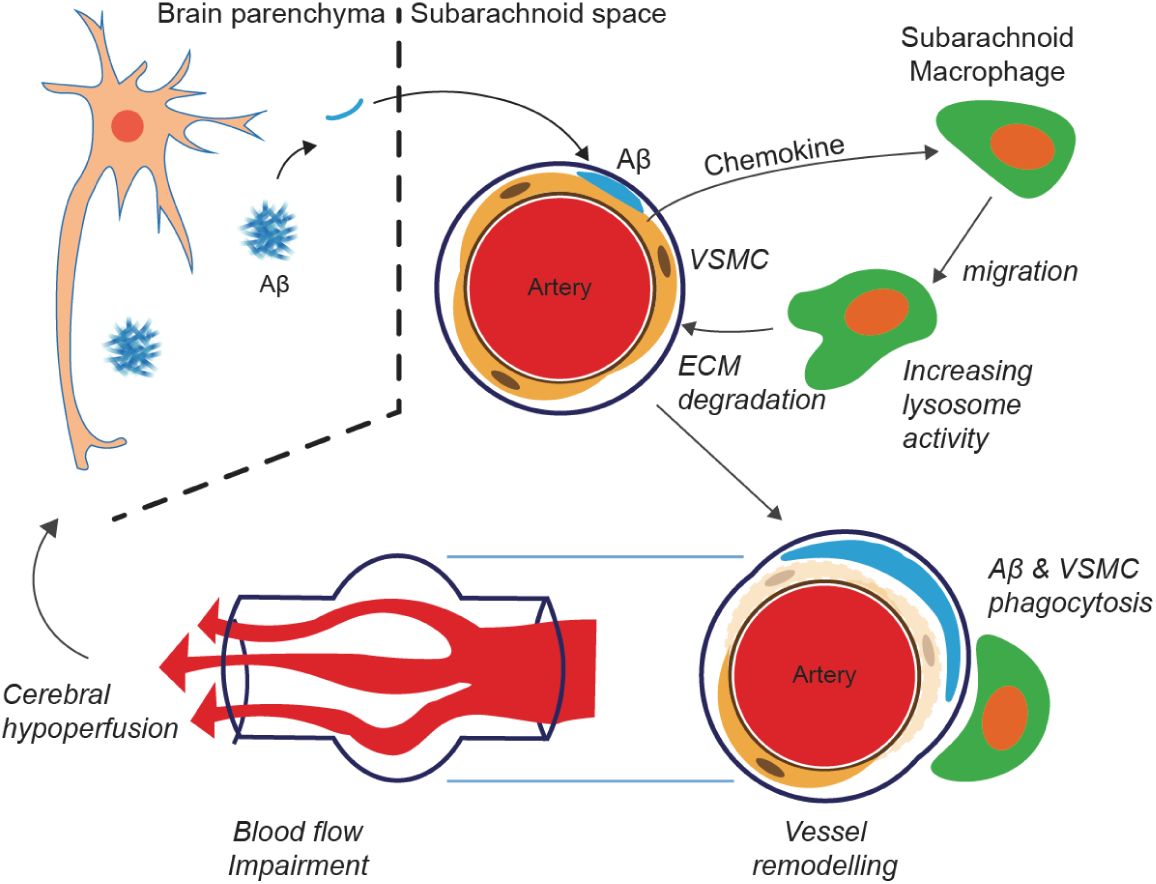
Schematic illustration of subarachnoid macrophage involvement in Aβ-associated cerebral vascular remodeling and hypoperfusion. Aβ originating from the brain parenchyma accumulates around the arteries in the subarachnoid space, triggering the release of chemokines from vascular smooth muscle cells (VSMCs). This promotes the migration of subarachnoid macrophages toward the affected vessel. Activated macrophages enhance lysosomal activity and contribute to the degradation of the extracellular matrix (ECM), leading to vessel remodeling. These changes impair blood flow and result in cerebral hypoperfusion, which may further exacerbate neurodegenerative processes in Alzheimer’s disease.

## METHODS

### KEY RESOURCES TABLE

| REAGENT or RESOURCE | SOURCE | IDENTIFIER |
| --- | --- | --- |
| **Antibodies** |  |  |
| Mouse MMR/CD206 Antibody | R&D Systems | Cat# AF2535; RRID: AB_2063012 |
| Mouse Anti-Aβ17-24 antibody (clone 4G8) | Biolegend | Cat# SIG-39220; RRID: AB_662812 |
| Mouse Anti-β-Amyloid (1–16, clone 6E10) | Biolegend | Cat# 803014; RRID: AB_662798 |
| Rabbit Anti β-Amyloid (D54D2) XP® Rabbit mAb | Cell Signaling Technology | Cat# 8243; RRID: AB_2797642 |
| Rat FITC-CD169 (Siglec-1) Antibody | Thermo Fisher Scientific | Cat# MA5-28187; RRID: AB_2745161 |
| Mouse Alexa Fluor 647-anti-α smooth muscle actin (1A4/asm-1) | Novus Biologicals | Cat# NBP2-34522AF647; RRID: AB_3287561 |
| Rabbit Anti α-Smooth Muscle Actin (D4K9N) XP® Rabbit mAb | Cell Signaling Technology | Cat# 19245; RRID: AB_2734735 |
| Rat LYVE1 Antibody | Thermo Fisher Scientific | Cat# 41-0443-82; RRID: AB_2573596 |
| Mouse Anti-NeuN Antibody | MilliporeSigma | Cat# MAB377; RRID: AB_2298772 |
| Donkey Anti-Rabbit IgG H&L Alexa Fluor 488 | Abcam | Cat# ab150073; RRID: AB_2630356 |
| Donkey Anti-Goat IgG H&L Alexa Fluor 594 | Abcam | Cat# ab150132; RRID: AB_2714037 |
| Goat Anti-Mouse IgG H&L Alexa Fluor 555 | Thermo Fisher Scientific | Cat# A21425; RRID: AB_141780 |
| Goat Anti-Rat IgG H&L Alexa Fluor 555 | Thermo Fisher Scientific | Cat# A21434; RRID: AB_2535850 |
| Goat Anti-Rabbit Alexa Fluor 594 – FluoroNanogold™ | (Thermo Fisher / Nanoprobe) | Cat# 7303; RRID: AB_3713462 |
| Mouse APC/Cyanine7 anti-mouse CD45.2 | Biolegend | Cat# 109824; RRID: AB_312978 |
| Rat BV605 anti-Ly6C | BD Biosciences | Cat# 563011; RRID: AB_2744479 |
| Rat BV711 anti-Ly6G | BioLegend | Cat# 127643; RRID: AB_2562265 |
| Rat APC-CD206 | Biolegend | Cat# 141707; RRID: AB_2573609 |
| Rat CD11b-PE | Invitrogen | Cat# 12-0112-82; RRID: AB_1089903 |
| Rat Alexa Fluor®594 anti-mouse CD45 | Biolegend | Cat# 103144; RRID: AB_312948 |
| Rat FITC anti-mouse CD11b | Biolegend | Cat# 101206; RRID: AB_312977 |
| Rat PE anti-mouse CD206 | Biolegend | Cat# 141705; RRID: AB_10896421 |
| Rat PE-Cy7 anti-mouse TMEM119 (clone V3RT1GOsz) | Invitrogen | Cat# 25611982; RRID: AB_2848312 |
| Goat Anti Iba1 | Abcam | Cat# ab5076; RRID: AB_2224402 |
| Mouse Anti-Rat IgG2a - Alexa Fluor 488 | Biolegend | Cat# 407514; RRID: AB_2910464 |
| Donkey Anti-Rabbit - Alexa Fluor 647 | Abcam | Cat# ab150075; RRID: AB_2752244 |
| Donkey Anti-Mouse - Alexa Fluor 405 | Abcam | Cat# ab175658; RRID: AB_2687445 |
| Rabbit Anti-MRC1 antibody | Sigma-Aldrich | Cat# HPA004114; RRID: AB_1846270 |
| TruStain FcX™ (anti-mouse CD16/32) Fc Block Antibody | Biolegend | Cat#101320<br>RRID: AB_1574975 |
| Anti-Mouse CD16/CD32 (Mouse BD Fc Block™) | BD Biosciences | Cat#553141; RRID: AB_394656 |
| **Biological samples** |  |  |
| Human post-mortem brain tissue (occipital lobe, CAA patients) | Brain Research Institute, Niigata University, Niigata, Japan | Two patients: 87 y/o Thal stage 1, 83 y/o Thal stage 2; Formalin-fixed; Ethics approval: Nagoya University 2025-0154; Informed consent obtained from legally authorized representatives |
| **Chemicals, peptides, and recombinant proteins** |  |  |
| Ketamine | Daiichi Sankyo Healthcare | Cat#S9019698 |
| Xylazine | Elanco | Cat#E136566 |
| Methoxy-X04 | Tocris Bioscience | Cat#4920/10 |
| Dextran, Texas Red™, 70,000 MW, neutral | Thermo Fisher Scientific / Invitrogen™ | Cat#D1830 |
| D-PBS(-) without Ca and Mg, liquid | Nacalai Tesque | Cat#14249-24 |
| 4% paraformaldehyde | Nacalai Tesque | Cat#09154-56 |
| OCT compound | Sakura Finetek | Cat#4583 |
| Tomato Lectin–DyLight 649 | Thermo Fisher Scientific | Cat#L32472 |
| Vectashield Antifade Mounting Medium with DAPI | Vector Laboratories | Cat# H1800 |
| Osmium tetroxide | Nacalai Tesque | Cat#2572701 |
| Sodium cacodylate buffer | Nacalai Tesque | Cat#0651664 |
| Potassium ferrocyanide | Katayama | Cat#244580 |
| CaCl<sub>2</sub> | Sigma | Cat#10043524 |
| Ethanol | Sigma | Cat#0907704 |
| Propylene oxide | Wako | Cat#16505026 |
| Quetol 812 resin | Electron Microscopy Sciences | Cat#14900 |
| Uranyl acetate | Nagoya University | N/A |
| Lead citrate | Wako | Cat#12101722 |
| Hank's Balanced Salt Solution | Gibco, Thermo Fisher Scientific | Cat#14025092 |
| DNAseI | Roche | Cat#11284932001 |
| Trypsin/EDTA Solution, with Phenol Red | Nacalai Tesque | Cat#32777-15 |
| Fetal Bovine Serum, qualified, Brazil | Gibco, Thermo Fisher Scientific | Cat#10270106 |
| Percoll Plus® | Cytiva | Cat#17-5445-01 |
| Red Blood Cell Lysis Buffer | Sigma-Aldrich | Cat#11814389001 |
| Fixable Viability Dye eFluor™ 506 | Invitrogen, Thermo Fisher Scientific | Cat#65-0866-14 |
| QIAzol Lysis Reagent | Qiagen | Cat#79306 |
| 2-mercaptoethanol | Nacalai Tesque | Cat#21417-52 |
| Collagenase NB 8 | Collagenase NB 8 | Cat#S1745602 |
| HEPES | Nacalai Tesque | Cat#17514-15 |
| Norland Optical Adhesive-61 | Norland Products Inc. | Cat#37-322 |
| G-CEM ONE adhesive enhancing primer | GC Corporation | Cat#228AKBZX00104000 |
| HiLyte Fluor 555 Labeling Kit -NH<sub>2</sub> | Dojindo Molecular Technologies, Inc. | Cat# LK14 |
| PLX-3397 (Pexidartinib) | MedChemExpress | Cat#HY-16749 |
| **Critical commercial assays** |  |  |
| Chromium Next GEM Single Cell 3' Reagent Kit v3.1 | 10x Genomics | Cat# 1000128 |
| SMART-Seq Stranded Kit | Clontech Laboratories | Cat# 634442 |
| **Deposited data** |  |  |
| Bulk RNA-sequencing data | This paper | DDBJ Database: PRJDB37973 |
| Single cell RNA-sequencing data | This paper | DDBJ Database: PRJDB37973 |
| **Experimental models: Organisms/strains** |  |  |
| Mouse *App<sup>NL-G-F/NL-G-F</sup>* | Model developed by Dr. T. Saito *et al.*<sup>28</sup> | Strain #: C57BL/6-*App<sup>tm3(NL-G-F)</sup>Tcs*>/TcsRbrc; RRID: RBRC06344 |
| Mouse *Cx3cr1<sup>GFP/GFP</sup>* | The Jackson Laboratory | Strain #:005582; RRID: IMSR_JAX:005582 |
| Mouse *C57BL/6* | CLEA Japan, Inc | C57BL/6JJcl; IMSR_JCL:JCL:MIN-0003 |
| **Software and algorithms** |  |  |
| ImageJ/Fiji v2.16.0 | Schindelin *et al.*<sup>42</sup> | RRID: SCR_002285; https://imagej.net/software/fiji/downloads |
| Illumina Casava v1.8.2 | Illumina | RRID: SCR_001802; http://support.illumina.com/sequencing/sequencing_software/casava.html |
| TopHat v2.0.13 | Johns Hopkins University | RRID: SCR_013035; https://ccb.jhu.edu/software/tophat/index.shtml |
| Bowtie2 v2.2.3 | Bowtie2 project | RRID: SCR_016368; https://bowtie-bio.sourceforge.net/bowtie2/index.shtml |
| SAMtools v0.1.19 | SAMtools project | RRID: SCR_002105; http://htslib.org/ |
| R v4.4.3 | R Project for Statistical Computing | RRID: SCR_001905; https://www.R-project.org |
| DAVID | NIH DAVID Bioinformatics | RRID: SCR_001881; https://davidbioinformatics.nih.gov/ |
| GraphPad Prism v10.4.1 | GraphPad Software | RRID: SCR_002798; https://www.graphpad.com/ |
| IMARIS v9.8.1 | Bitplane / Oxford Instrument | RRID: SCR_007370; https://imaris.oxinst.com/ |
| Line-Scan Particle Image Velocimetry (LS-PIV) | Kim *et al.*<sup>44</sup> | RRID: N/A; https://doi.org/10.1371/journal.pone.0038590 |
| Cell Ranger v9.0.1 | 10x Genomics | RRID: SCR_017344; https://www.10xgenomics.com/support/software/cell-ranger/downloads |
| Seurat package v5.2.1 | Satija Lab / R package | RRID: SCR_016341; https://cran.r-project.org/package=Seurat |
| GSEA (Gene Set Enrichment Analysis) v4.3.3 | Broad Institute | RRID: SCR_003199; https://www.broadinstitute.org/gsea/ |
| **Other** |  |  |
| Double-layered coverslip (3.5 and 4.5 mm) | Matsunami, Osaka, Japan | Custom made |
| Two-photon microscope | NIKON | A1R MP+ |
| Confocal fluorescence microscope | Olympus, Japan | FV3000 |
| Transmission electron microscope | JEOL Ltd, Japan | JEM-1400 |
| Cell sorter | Becton, Dickinson and Company | BD FACSMelody™ |
| Cell sorter | Sony Biotechnology, Japan | Sony MA900 |
| Laser speckle contrast imaging system | RWD Life Science, China | RFLSI III |
| Illumina HiSeq 6000 sequencing platform | Illumina, USA | HiSeq 6000 |
| Illumina NovaSeq 6000 sequencing platform | Illumina, USA | NovaSeq 6000 |
| Ultramicrotome | Leica Microsystems, Germany | EM UC7 |
| Fluorescence microscope | Keyence corporation | BZ-X800 |
| Light-Sheet Microscope | Miltenyi Biotec | UltraMicroscope Blaze Light-Sheet Microscope |

### EXPERIMENTAL MODEL AND STUDY PARTICIPANT DETAILS

#### Human brain samples

Post-mortem brain samples were obtained from two patients diagnosed with CAA, with Thal stage 1 (87 years old) and Thal stage 2 (83 years old). Formalin-fixed blocks were sampled from the occipital lobes and underwent a tissue-clearing protocol. First, delipidation was performed using 10% 1,2-hexanediol at 45°C for 72 h, followed by bleaching with 1% hydrogen peroxide at 45°C for 48 h. Cleared samples were stained with anti-Aβ_17-24_ antibody (1:100; BioLegend, SIG-39220) labeled using the HiLyte Fluor 555 Labeling Kit -NH₂ (Dojindo Molecular Technologies, Inc.), and Alexa Fluor 647-conjugated anti-α smooth muscle actin (SMA) antibody (1:100; Novus Biologicals, 1A4/asm-1) for 7 days. Samples were then dehydrated using methanol and optically cleared using benzyl alcohol and benzyl benzoate (BABB). The tissue was imaged using the Ultra Microscope Blaze light-sheet scanning microscope. Raw images were processed and analyzed using Imaris software (Bitplane, USA).

This study was approved by the Institutional Ethics Committee of Nagoya University (approval number: 2025-0154). Written informed consent for the use of post-mortem brain tissues was obtained from the legally authorized representatives of the donors, in accordance with the Declaration of Helsinki and relevant national guidelines.

#### Animals

All animal procedures were conducted in accordance with the Animal Care and Use Committee guidelines of Nagoya University Graduate School of Medicine. The experiments were conducted in accordance with the ARRIVE guidelines. All efforts were made to minimize the number of animals used and their suffering. Mice were housed in a 12-h light/dark cycle with free access to food and water. To model AD pathology, we used male *App^NL-G-F/NL-G-F^* mice that contain Swedish, Iberian, and Arctic mutations. To visualize macrophages *in vivo*, *App^NL-G-F/NL-G-F^* mice were crossbred with *Cx3cr1^GFP/GFP^* mice. We used *App^NL-G-F/NL-G-F^ Cx3cr1^GFP/+^* mice for *in vivo* imaging. Age-matched wild-type *C57BL/6* mice or *Cx3cr1^GFP/+^* were used as controls.

### METHOD DETAILS

#### Surgical preparation for *in vivo* imaging

Mice were anesthetized with an intraperitoneal injection of ketamine (74 mg/kg) and xylazine (10 mg/kg). After shaving and disinfecting the scalp with 70% ethanol, the skull was exposed. A craniotomy (4 × 4 mm) was performed over the somatosensory cortex while preserving the dura mater. A double-layered coverslip (3.5 and 4.5 mm; Matsunami, Osaka, Japan) was placed onto the craniotomy site and fixed using Norland Optical Adhesive-61 (Norland, Jamesburg, NJ, USA) under ultraviolet light.

To stabilize the head for imaging, a custom-made metal plate was attached to the skull using the G-CEM ONE adhesive enhancing primer and paste (GC Corporation, Tokyo, Japan). Throughout the procedure, body temperature was maintained using a heating pad. The mice were allowed to recover for at least 3 weeks before subsequent *in vivo* imaging experiments.

#### Two-photon imaging

*In vivo* two-photon imaging was performed using a NIKON A1 laser-scanning microscope equipped with 16× (NA 0.8) and 20× (NA 1.0) water-immersion objective lenses (Nikon Instech Co., Ltd., Tokyo, Japan). Imaging was performed using a Ti:sapphire laser (Chameleon Ultra II, Coherent, CA, USA) at excitation wavelengths of 820 and 950 nm. Emitted fluorescence signals were collected with non-descanned detectors. Image acquisition was performed using NIS-Elements software (Nikon Instech Co., Ltd.), and the data were exported as ND files. Image processing and analysis were conducted using ImageJ/Fiji (v2.16.0) ^42^.

To visualize Aβ deposition *in vivo*, methoxy-X04 (2 mg/kg body weight; Tocris, Bristol, UK) was administered intraperitoneally 1 day before imaging. For vascular visualization, 70 kDa Dextran-Texas Red (2.5% w/v in sterile saline; Thermo Fisher Scientific) was injected intravenously (50 µL via the retro-orbital vein) immediately prior to imaging.

To assess the progression of cortical Aβ deposition, longitudinal two-photon imaging was performed weekly from 24 to 36 weeks of age in *App^NL-G-F/NL-G-F^ Cx3cr1^GFP/+^* mice. Z-stack images were collected from the cortical surface to a depth of 100 µm, with 2-µm step intervals; the field of view was 2943.49 × 2943.49 µm (1.2872 pixels/µm). Imaging was performed in XYCZT format with a single time point (T = 1). Galvano scanning mode was used with a zoom factor of 1.0 and no line averaging.

To investigate dynamic interactions between SAMs and Aβ deposits, high-resolution time-lapse imaging was performed in 32- to 36-week-old mice. Images were captured over a 3-h period, with 3-min intervals. The imaging volume covered 265.17 × 265.17 µm in the XY plane and 50 µm in the Z direction, with 1-µm step intervals. Macrophage motility, including velocity and travel distance, was analyzed using the Manual Tracking plugin in Fiji (v2.16.0). To quantitatively assess macrophage morphology, we calculated cell circularity using the formula: Circularity = (4 × π × Area) / (Perimeter²). This metric ranges from 0 (highly irregular) to 1 (perfect circle). Area and perimeter were obtained from segmented fluorescence images using Fiji (v2.16.0), following thresholding and particle analysis on maximum intensity projections.

To evaluate vascular pulsatility, time-lapse imaging was conducted over a 60-s duration, capturing a single focal plane (103.89 × 103.89 µm) at 15 frames/s. Changes in vessel diameter were analyzed using the Vasometric plugin in Fiji ^43^.

Blood velocity measurements were obtained via line-scan two-photon microscopy, with scan lines oriented longitudinally along individual leptomeningeal vessels to align with the direction of blood flow. The resulting space-time (XT) images captured red blood cell movement as diagonal streaks, at a spatial resolution of 0.2994 µm/pixel and a temporal resolution of 0.9687 ms/line. Velocity quantification employed the Line-Scan Particle Image Velocimetry technique, which analyzes red blood cell displacement by cross-correlating XT image segments, enabling calculation of the mean red blood cell velocity over a 30-s acquisition period ^44^.

#### Immunohistochemistry

*App^NL-G-F^* mice were deeply anesthetized via intraperitoneal injection of ketamine (74 mg/kg) and xylazine (10 mg/kg), followed by transcardial perfusion with 20 mL of phosphate-buffered saline (PBS) and 20 mL of 4% paraformaldehyde (Nacalai Tesque, Japan). Brains were post-fixed overnight at 4°C and cryoprotected in 30% sucrose in PBS at 4°C until fully saturated. The brains were embedded in OCT compound (Sakura Finetek, USA) and stored at −20°C prior to sectioning.

Coronal sections (25 µm thickness) targeting the somatosensory cortex were prepared using a cryostat. Sections were washed in PBS for 5 min to remove residual OCT. Permeabilization was performed using 0.5% Triton X-100 in PBS for 10 min, followed by three washes (5 min each) in 0.05% Triton X-100 in PBS.

To block non-specific binding, sections were incubated in blocking buffer (PBS containing 5% donkey serum and 1% bovine serum albumin) for 1 h at room temperature. Primary antibodies diluted in blocking buffer were applied overnight at 4°C. The following antibodies were used: anti β-Amyloid (D54D2) XP Rabbit mAb (1:500; Cell Signaling Technology, 8243), anti-β-Amyloid (1–16, clone 6E10) antibody (1:500; BioLegend, 803014), mouse MMR/CD206 antibody (1:500; R&D Systems, AF2535), LYVE1 antibody (1:200; Thermo Fisher Scientific, 41-0443-82), FITC-CD169 (1:50; Siglec-1) antibody (Thermo Fisher Scientific, MA5-28187), anti-NeuN antibody (1:500; MilliporeSigma, MAB377), anti α-Smooth Muscle Actin (D4K9N) XP Rabbit mAb (1:500; Cell Signaling Technology, 19245), anti Iba1 (1:200; Abcam. Ab5076).

After washing (3 × 5 min in 0.05% Triton X-100 in PBS), sections were incubated for 1 h at room temperature in the dark with the following secondary antibodies: Donkey Anti-Rabbit IgG H&L (Alexa Fluor 488, Abcam, ab150073), Goat Anti-Mouse IgG H&L (Alexa Fluor 555, Thermo Fisher Scientific, A21425), Goat Anti-Rat IgG H&L (Alexa Fluor 555, Thermo Fisher Scientific, A21434), Donkey Anti-Goat IgG H&L (Alexa Fluor 594, Abcam, ab150132), Mouse Anti-Rat IgG2a (Alexa Fluor 488, Biolegend, 407514), Donkey Anti-Rabbit (Alexa Fluor 647, Abcam, Ab150075), and Donkey Anti-Mouse (Alexa Fluor 405, Abcam, Ab175658). After three additional PBS washes, the sections were stained with Tomato Lectin–DyLight 649 (1:200, Thermo Fisher Scientific, USA) for 30 min in the dark at room temperature. Nuclei were counterstained using Vectashield Antifade Mounting Medium with DAPI (Vector Laboratories, USA).

For human macrophage observation, cortical tissue from an 87-year-old individual with CAA was obtained post-mortem, formalin-fixed, and paraffin-embedded. Sections (10 µm) were deparaffinized, rehydrated, and first treated with formic acid for antigen retrieval, followed by heat-induced antigen retrieval in citrate buffer (pH 6.0). After permeabilization and blocking with donkey serum, sections were incubated overnight at 4°C with primary antibodies against Rabbit Anti-MRC1 (1:200, Sigma-Aldrich, HPA004114) and Mouse anti-β-Amyloid (1–16, clone 6E10) antibody (1:500; BioLegend, 803014), followed by Donkey Anti-Rabbit IgG H&L (Alexa Fluor 488, Abcam, ab150073) and Goat Anti-Mouse IgG H&L (Alexa Fluor 555, Thermo Fisher Scientific, A21425). Tomato lectin conjugated to Alexa Fluor 647 was applied after secondary incubation to label vessel walls, and nuclei were counterstained with DAPI. Sections were mounted with antifade medium and imaged using a confocal laser-scanning microscope.

*Z*-stack images (1 µm step size) were acquired using an Olympus FV3000 confocal fluorescence microscope with a 40× objective lens. Maximum intensity projections were generated using Fiji (v2.16.0).

#### TEM tissue preparation

*App^NL-G-F/NL-G-F^* mice (12 months old) were deeply anesthetized and perfused transcardially with 0.1 M phosphate buffer (pH 7.4) containing heparin, followed by 4% paraformaldehyde and 0.05% glutaraldehyde in 0.1 M sodium cacodylate buffer with 1 mM CaCl₂. Brains were dissected, cut into 3-mm blocks, and post-fixed overnight with gentle agitation. Vibratome sections (100 µm) were incubated with anti β-Amyloid (D54D2) XP Rabbit mAb (1:200; Cell Signaling Technology, 8243) in 1% BSA/PBS at 4°C overnight, followed by Goat Anti-Rabbit Alexa Fluor 594–FluoroNanogold antibody (1:100, Nanoprobes, 7303). Sections were post-fixed in 1% osmium tetroxide with 0.1% potassium ferrocyanide in sodium cacodylate buffer containing 1 mM CaCl₂ for 1 h at room temperature, rinsed, and dehydrated through a graded ethanol series followed by 100% propylene oxide. Samples were infiltrated with Quetol 812 resin via stepwise resin–solvent exchanges, degassed under vacuum, and embedded in fresh resin. Polymerization was performed at 60°C for 2–3 days. Tissue blocks were serially sectioned into ultrathin slices with a diamond knife on a Leica EM UC7 ultramicrotome, stained with uranyl acetate and lead citrate, and examined using a JEOL JEM-1400 transmission electron microscope.

#### Flow cytometry

Mice were euthanized via intraperitoneal injection of ketamine (74 mg/kg) and xylazine (10 mg/kg). The cerebral cortex, including subdural tissue, was rapidly harvested and placed into ice-cold Hank’s Balanced Salt Solution (HBSS) supplemented with 0.1 mg/mL DNase I (Roche, Switzerland). Tissues were minced using a sterile razor blade and enzymatically digested with 0.25% trypsin/EDTA (1 mmol/L; Nacalai Tesque, Japan) for 30 min at 4°C, followed by an additional 5 min at 37°C. The reaction was quenched with 10% fetal bovine serum (FBS) and washed with HBSS.

The resulting suspension was gently triturated using a 1,000-µL pipette and filtered through a 70-µm mesh to remove debris. Cells were separated from remaining tissue fragments using 36% Percoll™ Plus (Cytiva, Sweden), and red blood cells were eliminated using RBC lysis buffer (Roche, Germany). The final cell pellet was washed with PBS and resuspended in FACS buffer (2% FBS in PBS).

For staining, cells were first incubated with TruStain FcX (anti-mouse CD16/CD32; BioLegend, USA) at a 1:100 dilution for 15 min at 4°C to block Fc receptors. Subsequently, cells were stained with the following fluorochrome-conjugated antibodies (all at 1:100 dilution): Alexa Fluor 594 anti-mouse CD45 (BioLegend, 103144), FITC anti-mouse CD11b (BioLegend, 101206), PE anti-mouse CD206 (BioLegend, 141705), and PE-Cy7 anti-mouse Tmem119 (Invitrogen, 25611982). Staining was carried out for 15 min at 4°C in the dark.

Following staining and washing, cells were sorted using a BD FACSMelody™ Cell Sorter (BD Biosciences, USA). Dead cells were excluded by staining with Fixable Viability Dye eFluor™ 506 (Thermo Fisher Scientific, USA). Macrophages were identified as CD11b⁺/CD45^high^/Tmem119^low^/CD206^high^, as previously described ^45^. Target cell populations were sorted into QIAzol Lysis Reagent (Qiagen Sciences, USA) supplemented with 1% 2-mercaptoethanol (Nacalai Tesque, Japan) for downstream RNA extraction.

#### Bulk RNA sequencing

Single-cell suspensions were obtained from 7-month-old *App^NL-G-F^* mice and age-matched *C57BL/6* control mice for bulk RNA sequencing. Total RNA was extracted using the SMART-Seq Stranded Kit (Clontech Laboratories), and cDNA libraries were prepared according to the manufacturer’s instructions. Sequencing was performed using paired-end reads on the Illumina HiSeq 6000 platform (Illumina). Base calling was carried out using Illumina Casava software (v1.8.2).

Raw sequencing reads were aligned to the mouse reference genome (mm10) using TopHat (v2.0.13) in conjunction with Bowtie2 (v2.2.3). SAMtools (v0.1.19) was used for subsequent processing and indexing of the aligned reads. Differential gene expression analysis was conducted, and significantly regulated genes were identified using an adjusted p-value <0.05 and a fold-change threshold of ≥1.5. These genes were subjected to Gene Ontology and pathway enrichment analysis using DAVID ^46,47^, followed by Gene Set Enrichment Analysis (GSEA) ^48,49^.

#### Macrophage isolation and sorting for scRNA-seq

We collected the brain cortex, including the brain surface, and minced it using a razor to get 1 mm^3^ cuts. Brains were enzymatically dissociated by incubation in 5 mL of enzyme solution (1× HBSS (+), 10 mM HEPES, 10% FBS) containing 12.5 mg Collagenase NB 8 (S1745602, Nordmark) per tissue at 37°C for 15 min. The tissue was gently triturated with a 1-mL pipette, incubated for an additional 15 min, then homogenized through an 18-G needle and incubated for an additional 15 min. The suspension was then passed through a 20-G needle, treated with 0.5 M EDTA (100 µL), and centrifuged at 500 × g for 5 min at 4°C. The pellet was resuspended in 25% Percoll and layered over 65% Percoll for density gradient centrifugation (1000 × g, 20 min, 4°C, low acceleration, no brake). The interface was collected and washed once with PBS containing 2% FBS and 10-mM EDTA. Fc receptors were blocked with anti-CD16/32 (Fc block, 2.4G2, BD Biosciences) for 10 min at 4°C prior to antibody staining.

Cells were stained with the following antibodies: APC/Cyanine7 anti-mouse CD45.2 (109824, BioLegend), Ly6C-BV605 (563011, BD Biosciences), BV711 anti-mouse Ly6G (127643, BioLegend), CD206-APC (141707, BioLegend), and CD11b-PE (M1/70, 12-0112-82, Invitrogen). CAMs were defined as CD45⁺ CD11b⁺ Ly6C⁻ Ly6G⁻ CD206⁺ cells, and microglia were defined as CD45⁺ CD11b⁺ Ly6C⁻ Ly6G⁻ CD206⁻ cells. The stained cells were sorted using a Sony MA900 cell sorter (Sony Biotechnology, Japan) and collected into tubes containing FACS buffer (PBS + 2% FBS) without EDTA.

#### scRNA-seq and data analysis

Sorted CAMs and microglia were subjected to scRNA-seq using the Chromium Next GEM Single Cell 3′ Reagent Kit v3.1 (10x Genomics), targeting ∼10,000 cells per sample. Libraries were sequenced on an Illumina NovaSeq 6000 platform. Raw data were processed using Cell Ranger (v 9.0.1, 10x Genomics) with the GRCm39 reference genome. The infrastructure of the Omics Science Center’s Secure Information Analysis System, located at the Medical Institute of Bioregulation at Kyushu University, provided computational resources. Downstream analyses were performed in R (v4.4.3) using the Seurat package (v5.2.1). For quality control, cells with more than 5% mitochondrial counts and extreme unique feature counts (either over 4,500 or under 200) were removed. Data were normalized with the NormalizeData()function. The data were then combined using FindIntegrationAnchors() and IntegrateData() functions. Principal component analysis (PCA) was performed using the combined data. A scree plot was generated using the ElbowPlot() function to select the principal components (PCs) by identifying the last PC at which the explained variance reaches a plateau. Selected PCs were used to calculate nearest-neighbor distances and to perform Louvain clustering using the FindNeighbors() and FindClusters() functions. Uniform manifold approximation and projection were then used to visualize clusters. Differentially expressed genes were identified using the Wilcoxon rank-sum test with Benjamini–Hochberg correction (adjusted p value <0.01). Functional annotation and enrichment analyses were performed using DAVID (https://david.ncifcrf.gov/) for Gene Ontology and Kyoto Encyclopedia of Genes and Genomes pathway terms.

#### Arterial hemodynamic parameters

Cerebral blood perfusion was assessed in 9-month-old *App^NL-G-F/NL-G-F^ Cx3cr1^GFP/+^* mice and *Cx3cr1^GFP/+^* controls. A craniotomy and headplate fixation were performed 3 weeks prior to imaging. Mice were imaged using a two-photon microscope at an excitation wavelength of 820 nm to localize Aβ-associated blood vessels.

Methoxy-X04, a fluorescent probe for Aβ, was administered intraperitoneally 1 day prior to *in vivo* imaging. To visualize the cerebrovascular network, 70 kDa Texas Red-conjugated dextran (Thermo Fisher Scientific) was injected intravenously. Blood flow in the target vessels was recorded using the RFLSI III laser speckle contrast imaging system (RWD Life Science, China). Mice were head-restrained using the fixed metal plate and allowed to habituate for 3 min before imaging. Acquired images were processed using RWD’s proprietary analysis software.

### QUANTIFICATION AND STATISTICAL ANALYSIS

All statistical analyses were performed using GraphPad prism version 10.4.1 (GraphPad software, USA). Data are expressed as mean ± standard deviation (SD), unless otherwise noted. For comparisons between two groups, unpaired or paired two-tailed student’s t-tests were used for normally distributed data, while the Mann–Whitney u test was used for non-parametric data. For analyses involving more than two groups, the one-way or two-way ANOVA followed by Dunnett’s, Tukey’s, or Sidak’s multiple-comparisons test was applied, where appropriate. For multiple group comparisons with non-parametric distributions, we used the Kruskal–Wallis test. Normality was assessed using the Shapiro–Wilk test. Correlations were assessed using Pearson’s correlation coefficient. Statistical significance was set at p < 0.05 or p < 0.01.

