## Supplemental Information for "Vicious circle of amyloid and leptomeningeal macrophages evokes vascular dysfunction in CAA"

**The PDF file includes:**

Videos S1 and S2

**Video S1. Time-lapse imaging of ring-shaped formation of Aβ deposition around the leptomeningeal arteries.**

Long-term imaging of the ring-shaped formation of Aβ deposition around the leptomeningeal arteries. The numbers in the top-right corner represent weeks. Scale, 25 μm. This video corresponds to Figure 1M.

**Video S2. Immune reaction of SAMs against Aβ deposition.**

Immune reaction between macrophages (green) and Aβ (blue) on leptomeningeal arteries (red) was tracked *in vivo* over time. The numbers in the top right represent hours and minutes. Scale, 50 μm. This video corresponds to Figure 2F.
